# Facing West Africa’s rosewood crisis: what determines the success of preserving and propagating the endangered barwood (*Pterocarpus erinaceus*)? A multi-model test of site and maternal-origin effects

**DOI:** 10.64898/2026.10.01.755929

**Authors:** Beda Innocent Adji

## Abstract

*Pterocarpus erinaceus* Poir. (Fabaceae), the barwood or vène, is at the heart of one of the most severe conservation crises currently affecting tropical timber species. It has become the most heavily traded timber species in the world by volume, at the centre of an illegal rosewood trade to Asia that has driven the species to near-total disappearance in several countries despite its listing on CITES Appendix II since 2016. Faced with this urgency, ex situ conservation and restoration programmes often rest on the untested assumption that selecting seed trees by size improves the quality of the offspring produced in the nursery. This study rigorously tests this hypothesis against the competing one of a determinism dominated by the planting site, using six complementary analytical approaches (continuous regression, linear mixed models with a site × seed-tree interaction, seed-mass mediation analysis, non-linear growth modelling of height and diameter, between-site phenotypic plasticity, and multi-predictor allometric equations), across two contrasting bioclimatic sites in Côte d’Ivoire (Korhogo: dry zone; Daloa: humid zone) and six seed-tree categories. Site exerts a decisive influence on growth at all ages (ANOVA, *p*<0.001 for height, diameter and biomass), whereas germination does not differ significantly between sites (70.3% at Daloa versus 61.6% at Korhogo; χ^*2*^=1.95; *p*=0.162). Seed-tree DBH does not affect growth in any of the tests (continuous regression, *p*>0.30; mixed model, *p*>0.07), and the seed-mass mediation analysis remains inconclusive, seed mass varying only slightly among seed trees in this species (0.07-0.08 g). The exponential growth model provides the best fit for height and diameter at both sites, and two-predictor allometric equations explain 96.2% of biomass variance compared with 89.9% for the single-predictor model. These results indicate that ex situ conservation and compensatory reforestation programmes for this endangered species would benefit from prioritising the choice of planting site over the dendrometric selection of seed trees. A recommendation directly applicable to conservation nurseries established in response to the rosewood crisis.

**Highlights:**

- Planting site strongly outweighs seed-tree size in juvenile growth;
- Site effects intensified from six to 24 months;
- Seed-tree size did not consistently predict offspring performance;
- Juvenile growth followed exponential trajectories across contrasting sites;
- Non-destructive allometry explained 96.2% of biomass variation.

## Introduction

Illegal international trade in threatened species is among the leading threats to tropical biodiversity, and West African rosewood constitutes one of the most emblematic cases of the past decade (Dumenu, 2019; Environmental, 2022). *Pterocarpus erinaceus* Poir. (Fabaceae), commonly known as barwood, vène or kosso, has become the most heavily traded timber species in the world by volume, driven by booming Asian demand for antique-style furniture « hongmu » (CITES, 2016). Chinese log imports of the species increased more than 2000-fold between the third quarter of 2009 and the third quarter of 2015, rising from 70 to approximately 149,000 m_3_ per quarter (CITES and CoP17 Prop, 2016). Over the 2015-2016 period alone, China imported an average of 764,000 m_3_ of West African rosewood per year, worth USD 840 million, representing the harvest of more than one million trees (CITES, 2016; CITES and CoP17 Prop, 2016). This pressure has driven the collapse of entire populations; the species is considered to have nearly disappeared from some countries, such as the Gambia, since 2011, despite multiple national logging bans, its Endangered (EN) status on the IUCN Red List, and its listing on CITES Appendix II since 2016 (Barstow, 2017; IUCN, 2018). Illegal trade nonetheless continues on a large scale despite these measures, as evidenced by discrepancies of several million tonnes between declared export and import volumes over the 2016-2020 period (Environmental, 2022; Environmental inverstigation agency, 2025).

Faced with this conservation crisis, nursery seedling production (for the restoration of degraded stands, the establishment of conservation seed orchards, or the compensatory reforestation programmes already mandated in some exporting countries (CITES, 2018)), constitutes a direct, immediately actionable lever, unlike international trade-control measures whose effectiveness remains largely compromised. This pathway is all the more necessary given that the species’ natural regeneration, like that of many Sahelian and Sudanian woody species, is itself compromised by recurrent bushfires and land-use change, with growth performance varying widely across ecological zones and soil properties (Asigbaase et al., 2024). Yet the success of these ex situ conservation programmes depends on practical choices rarely underpinned by scientific evidence: should seeds be sourced from the largest remaining seed trees, on the assumption that their offspring will be of higher quality, or does the choice of planting site matter more than the origin of the planting material? This question, far from anecdotal, directly determines the allocation of scarce resources in conservation programmes with limited means.

Adji et al. (2021), using a design comparing three contrasting environments (Korhogo, Daloa, and a controlled greenhouse in Montpellier), directly tested this question in Pterocarpus erinaceus and showed that the environment, but not the characteristics of the six sampled seed trees, significantly influenced germination parameters (*P*<0.05 for environment, *P*>0.05 for seed tree). Seed trees affected only leaf number and leaf length in seedlings, with no effect on height, diameter or leaf width. This result, obtained through a categorical analysis of variance limited to the 4-month stage, calls for confirmation using statistical methods of greater power and over a longer developmental window; precisely the aim of the present study. Tiika et al. (2019), in Ghana, examined the effect of mother-tree size, source and fruit-harvest timing on the quality and germinability of *Pterocarpus erinaceus* seeds, likewise concluding that there was no correlation between seed mass and germination percentage according to tree size. Conversely, Ansah et al. (2023) showed, in the same species in Ghana, that the maternal environment (temperature, soil nutrients) significantly influences the characteristics of the seeds produced, with repercussions for their germination potential; suggesting that the effect of the mother population’s environment might outweigh that of individual seed-tree size. More recently, work conducted in Togo and Côte d’Ivoire has characterised germination and growth of the species in nursery conditions (Nathalie et al., 2020) and tested various pretreatments and substrates to improve performance (Fulgence et al., 2024), without, however, isolating the effect of seed-tree size from that of the site with the methodological rigour required.

These results, although convergent, remain incomplete and methodologically limited. They call for a study simultaneously testing, within a single experimental design and using statistical tools of adequate power, the effect of individual seed-tree dendrometric characteristics and that of the planting site. The present study formulates three research questions: (i) does the planting site influence the germination and juvenile growth of Pterocarpus erinaceus? (ii) do the diameter and height of the original seed tree have an effect, whether direct or mediated by seed mass, on the performance of its offspring? and (iii) is this effect, or its absence, consistent with the pattern reported by Adji et al. (2021) at the 4-month stage, or does it change with seedling age? Three hypotheses are tested accordingly: H1, the site exerts a significant effect on growth, consistent with the bioclimatic contrast of the design; H2, seed-tree characteristics have no detectable direct effect on offspring performance, consistent with Tiika et al. (2019); H3, if an indirect effect exists via seed mass, its detection will be limited by the low intraspecific variability of this trait in this species. Six complementary analytical approaches are used to test these hypotheses: continuous regression, linear mixed models with a site × seed-tree interaction, seed-mass mediation analysis, non-linear modelling of growth trajectories (height and diameter), between-site phenotypic plasticity indices, and multi-predictor allometric equations compared by *AIC* (Akaike, 1974). By clarifying this determinism, this study aims to provide managers of ex situ conservation and compensatory reforestation programmes with a directly applicable scientific basis: faced with the accelerating scarcity of wild stands of the species, should these programmes focus their limited efforts on selecting large seed trees among the few remaining mature individuals, or on the judicious choice of planting site?

## Materials and methods

### Study sites

The study was conducted between April 2019 and July 2025, with several repetitions, in two experimental nurseries in Côte d’Ivoire, selected for their marked bioclimatic contrast along a north-south climatic gradient. The Korhogo forestry research station (9°57’0.556” N; 5°54’2.889” W), managed by the Centre National de Recherche Agronomique (CNRA), is representative of the Sudanian zone: dry tropical climate, annual rainfall of 817 to 1216 mm, temperatures of 26.6 to 35.7 °C, shallow, gravelly ferruginous and ferralitic soils, low in organic matter and strongly desaturated (Adji et al., 2023). The Jean Lorougnon Guédé University in Daloa (6°9’6.363” N; 6°43’8.1157” W), in the west-central part of the country, is representative of the sub-equatorial dense humid forest zone: rainfall of 1000 to 1900 mm/year, temperatures of 21 to 34 °C, deep, acidic ferralitic soils, desaturated in exchangeable bases but rich in organic matter (Adji et al., 2023; Akaffou et al., 2021). This pedoclimatic contrast makes it possible to test the site effect independently of the origin of the plant material, which is identical at both sites.

### Study species and plant material

*Pterocarpus erinaceus* Poir. (Fabaceae) is a tree that can reach 12 to 15 m in height (exceptionally 25 m), with a sparsely branched crown, characteristic of Sudano-Guinean savannas and open woodlands (Louppe and Ouattara, 1996; Segla et al., 2016). Despite multiple national logging bans across several African countries, it remains overexploited for multiple uses, and its illegal trade to Asia has led to its listing on CITES Appendix II in 2016, with an Endangered (EN) status on the IUCN Red List (CITES and CoP17 Prop, 2016; Dumenu, 2019; IUCN, 2018). Seeds were collected from 120 seed trees over several harvest campaigns (April and May 2019; April 2020 and May 2023) in four localities arranged along a bioecological gradient in Côte d’Ivoire: Katiola (5°7’35.814” W; 8°13’53.94” N), Niakara (5°18’40.73544” W; 8°40’47.97912” N), Korhogo (5°36’12.39612” W; 9°33’24.68988” N) and Sinématiali (5°22’59.999” W; 9°34’59.999” N). Seed trees were grouped into six distinct categories according to their dendrometric size. Six reference trees were selected at the CNRA forestry research station in Korhogo to represent a gradient of contrasting dendrometric characteristics (diameter at breast height and total height), and 19 additional seed trees with characteristics close to the reference tree of each group were then sought across all four provenance localities mentioned above, forming six groups of 20 seed trees (Table 1). Seed trees from the same locality were approximately 300 m apart from one another.

**Table 1.**
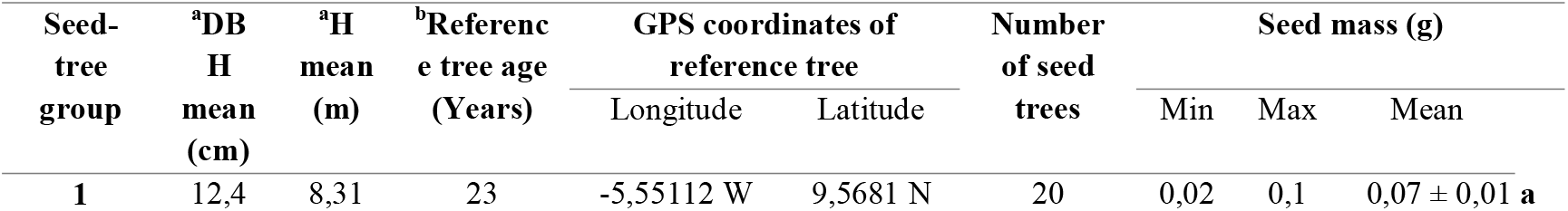
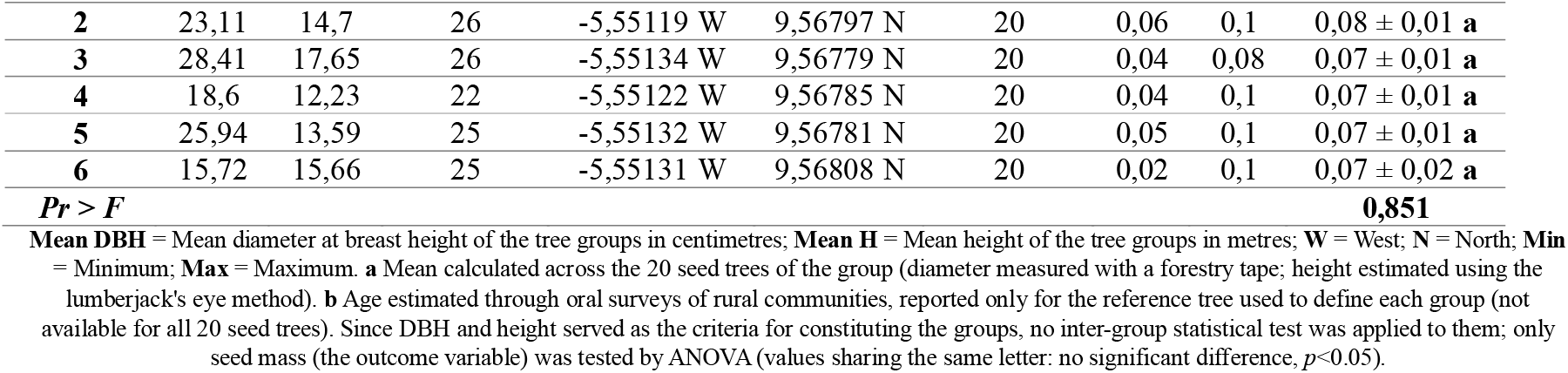
Dendrometric characteristics of the six seed-tree groups of *Pterocarpus erinaceus*.

The mean DBH and mean height of each group, calculated across the 20 seed trees, range respectively from 12.4 to 28.4 cm and from 8.3 to 17.7 m (Figure 1); it should be noted that these ranges partially overlap between certain groups, which retrospectively justifies the use, in the analyses presented, of continuous regression and mixed models treating DBH as a real variable rather than a simple categorical comparison, the latter approach being more sensitive to partial overlap between classes. The age of the reference tree in each group, by contrast, was estimated through oral surveys of rural communities and technicians at the CNRA forestry research station. This information is therefore available only for this single representative tree, not for all 20 seed trees in the group. Diameter was measured with a forestry tape and height was estimated using the lumberjack’s eye method. The plant material used is the property of rural populations; authorisation to use it for this study was given orally by the orchard owners.

**Figure 1.**
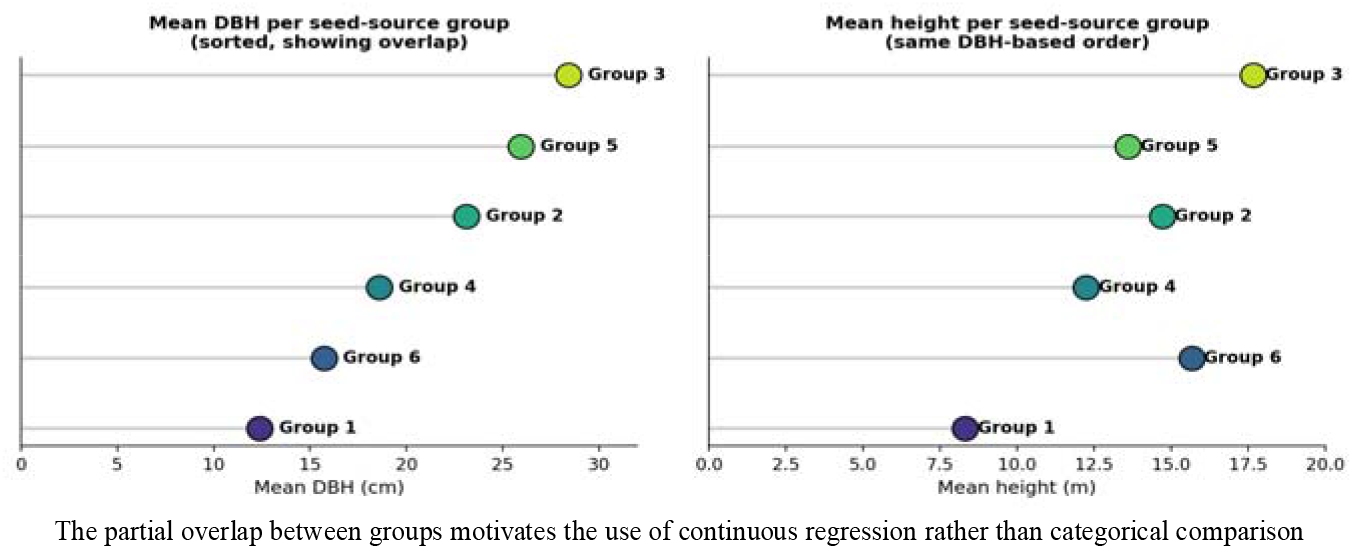
Dendrometric gradient (mean DBH and height) among the six seed-tree groups of *Pterocarpus erinaceus*, sorted by increasing DBH.

Seedlings and plants derived from the germination of seeds from the six seed-tree categories were monitored at different ages (4 months, 6 months, 1 year, 2 years, 5 years and 7 years). Seeds intended for germination and growth trials were extracted manually, sorted by flotation, and air-dried until moisture stabilised (4.5-5%), following the ISTA (2009). The present study specifically uses germination and growth data collected at 6, 12 and 24 months at the two contrasting sites of Korhogo and Daloa, constituting a subset of the broader experimental design described above; monitoring at 5 and 7 years, as well as the three other seed-collection localities (Katiola, Niakara and Sinématiali), are not covered by the analyses presented here (no provenance test).

### Experimental design

Seeds from each seed-tree group were divided into two lots destined respectively for Korhogo and Daloa. After soaking in tap water for a quarter of a day (6 hours) to break seed-coat dormancy, seeds were sown at a depth of 1.5-2 cm (with the lower end of the seed directed into the soil and the upper, wider end towards the surface) in polyethylene bags (20 × 10 cm) filled with local potting soil, at a rate of two seeds per bag, arranged in a block subdivided into six sub-blocks representing each seed-tree group. Each sub-block, containing 60 bags, was labelled with the serial number of the seed-tree group, i.e. 2 seeds × 60 bags × 6 groups × 2 sites = 1440 seeds for the trial. Seeds were collected and sown in subsequent years as experimental repetition and successive replacement. A phytosanitary treatment (granular FURADAN against rodents, DECIS against larval attacks) was applied, and maintenance consisted of daily watering and manual weeding.

### Germination monitoring

Germination, defined by the appearance of the hypocotyl at the substrate surface, was monitored daily (cumulative kinetics) at each site, allowing a cumulative germination model to be fitted and five classical parameters to be calculated (lag time, delay, rate, duration and germination percentage), following the definitions of Adji et al. (2023; 2020).

### Growth and biomass monitoring

After germination, 30 vigorous seedlings were randomly selected per seed-tree group and per site for non-destructive growth monitoring at 6, 12 and 24 months: total height (from collar to apex), collar diameter, number and dimensions of leaves, root length and diameter, measured using a graduated ruler, electronic callipers and ImageJ software for leaf area. A subsample of plants was harvested at the same ages for destructive biomass measurement (total, foliar, stem and root fresh and dry mass), after oven-drying (70 °C, 72 h) and weighing to the nearest 0.001 g, allowing allometric equations to be established.

### Data quality control

A systematic consistency check revealed, exclusively in the 12-month biomass measurement file, a substantial number of individuals (74 out of 145 with root data, i.e. 51%) exhibiting a root mass exceeding total plant mass, a physically impossible configuration. Further examination confirmed the origin of this anomaly: mean total dry biomass at 12 months was, at both sites, lower than that measured at 6 months; a growth regression that is itself physically impossible, indicating a likely systematic error in this specific file. Height and diameter data at 12 months, unaffected by this anomaly, were retained; only biomass variables at this age were excluded from the analyses. This anomaly, specific to the 12-month biomass measurement campaign, would warrant verification by the team that carried out this field data collection, for instance to rule out a balance calibration issue or a data-entry error occurring on that specific occasion. Detected outliers were treated as missing data rather than arbitrarily corrected, and explicitly documented. This step, rarely explicitly reported in the literature on the regeneration of African forest tree species, is presented here in the interest of reproducibility and scientific transparency.

### Analytical strategy

Rather than comparing seed-tree categories using classical analysis of variance, historically used in this species (Tiika et al., 2019), six complementary statistical approaches were used to maximise the power to detect any seed-tree effect and to clearly distinguish it from the site effect:

#### Continuous regression

Rather than comparing seed-tree groups against one another, as if tree size were a fixed category, we directly tested whether the actual diameter (DBH) and height of the seed tree can predict the performance of its offspring (much as one would test whether parents’ height predicts that of their children, by plotting a scatter plot and fitting a line through it). Specifically, each growth parameter was regressed on DBH and then on seed-tree height, separately at each age, by simple linear regression. For each growth parameter Y (height, collar diameter, total biomass) and at each measurement age, a simple linear regression model was fitted separately on the actual DBH and height of the original seed tree, rather than on a categorical group variable:

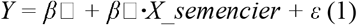

*Y is the growth parameter, X_seed-tree is the seed tree’s DBH or height, and β□ is the tested coefficient (fitted by ordinary least squares), or more explicitly, X_seed-tree alternately represents the seed tree’s DBH or height (continuous variable, in cm or m), β□ is the intercept, β□ is the tested regression coefficient (H0: β□ = 0), and ε is the residual error term, assumed i.i.d. ~ N(0, σ^2^). The model was fitted by ordinary least squares (OLS), separately for each parameter × age × predictor combination (18 regressions in total: 3 parameters × 3 ages × 2 predictors)*.

#### Linear mixed models

For each response *Y* (height, diameter, log-biomass), a linear mixed model was fitted with seed tree as a random effect (accounting for the non-independence of individuals originating from the same mother tree) and site, age and seed-tree DBH as fixed effects, including a DBH × site interaction term. Indeed, seedlings from the same seed tree are not entirely independent of one another: they share a common origin, much like siblings within the same family. Ignoring this relatedness can bias classical statistical tests. The mixed model corrects for this by treating each seed tree as a “family” whose own variability is estimated, while simultaneously testing the effect of DBH, site, and their interaction (is the effect of DBH different at Daloa and Korhogo?) on growth:

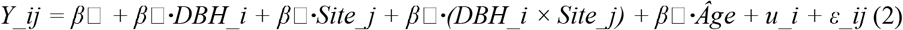

where *Y_ij* is the observed value for the individual from seed tree *i* at site *j*; *u_i* ~ *N*(0, σ^*2*^*_u*) is the random effect of seed tree *i* (accounting for between-seed-tree variance); ε*_ij* ~ *N*(0, σ^2^_ε) is the individual-level residual. Coefficient β□ tests the main (direct) effect of seed-tree DBH; β□ tests whether this effect differs by site (interaction). Models were fitted by restricted maximum likelihood (REML, Bates et al., 2015), using the lme4/lmerTest packages (*R*). For biomass, the response variable was log-transformed « log(total dry mass) » to stabilise variance, as the raw distribution was strongly right-skewed.

#### Mediation analysis and Sobel test

Even if seed-tree DBH has no direct effect on the growth of its offspring, it could have a hidden, indirect effect through an intermediate step: a large tree might produce heavier seeds, and these heavier seeds might in turn germinate better and produce more vigorous seedlings. We tested this pathway (DBH → seed mass → germination → growth) in several steps at the level of the six seed-tree groups (*n* = 6) through three simple sequential regressions (paths *a* to *c*), each testing one link in the chain (Baron and Kenny, 1986) :

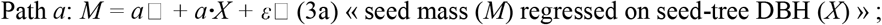

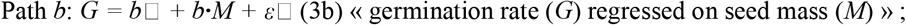

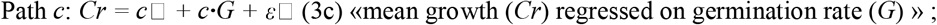

Total effect : *Cr = c’*□ *+ c’·X +* ε□ « growth regressed directly on seed-tree DBH, with no mediator », and the indirect effect of *X* on *G* via *M* is estimated by the product of the coefficients: Indirect effect = *a* × *b*. A complementary test (the Sobel test) allows verification of whether the overall indirect effect (through all links) is statistically significant, even if each link taken in isolation is not fully significant:

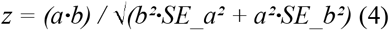

where *SE_a* and *SE_b* are the standard errors of coefficients *a* and *b* respectively. The *z* statistic is compared against the standard normal distribution (two-tailed test, α = 0.05 threshold).

#### Non-linear modelling of growth trajectories

Plants do not necessarily grow at a constant rate over time: some species grow slowly at first then accelerate, while others slow down as they age. We therefore compared three possible shapes of height-growth curve (a straight line, « steady growth »; a power-law curve, « progressively slowing growth »; and an exponential curve, « growth that accelerates over time ») to identify which best describes the actual seedling trajectory, separately for each site. The choice among these three curves was made using Akaike’s Information Criterion (*AIC*), an index that rewards the model offering the best trade-off between goodness of fit and simplicity (an unnecessarily complex model is penalised even if it fits the data slightly better). Three competing models were fitted to the individual height data (*H*) as a function of age (*t*, in months), separately by site, by non-linear least-squares regression (Levenberg-Marquardt):

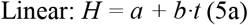

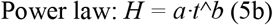

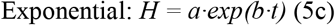

The three models were compared using Akaike’s Information Criterion, calculated from the residual sum of squares (*RSS*) on a common scale (raw, untransformed scale) to ensure a fair comparison between models:

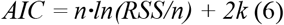

where *n* is the number of observations and *k* is the number of model parameters (*k* = 2 for the three models tested). The retained model is the one with the lowest *AIC* (ΔAIC > 2 being considered a substantial difference in support between models; Burnham and Anderson, 2002; Paine et al., 2012).

#### Phenotypic plasticity index

Beyond mean performance, we sought to determine whether certain seed trees produce offspring that are more « sensitive » to site conditions than others; that is, offspring whose growth varies markedly depending on whether they grow at Daloa or Korhogo (high plasticity), as opposed to offspring that behave similarly at both sites (low plasticity). A between-site phenotypic plasticity index of the RDPI type (*Relative Distance Plasticity Index*, Valladares et al., 2006) was calculated for height, by seed-tree group *i* and age *t*:

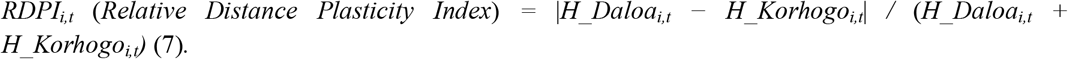

where *H_Daloa*_*i*,*t*_ et *H_Korhogo*_*i*,*t*_ are the mean heights of seed-tree group i at age t, at Daloa and Korhogo respectively. The index ranges from 0 (no plasticity, identical performance at both sites) to 1 (maximum plasticity, if performance differs greatly). The relationship between this index and seed-tree DBH was then tested by regression and by one-way analysis of variance (seed-tree group).

#### Allometric equations

Measuring the biomass (dry weight) of a plant requires cutting and oven-drying it; a destructive measurement, impossible to carry out across an entire nursery or reforestation trial. Allometric equations circumvent this problem: they relate biomass to simple, non-destructive measurements, such as collar diameter or height, taken on a sample of sacrificed plants, then applied to all other plants without having to cut them down. We compared a model using diameter alone with a model combining diameter and height, to determine which better predicts actual biomass. Total dry biomass (*B*) was modelled as a function of collar diameter (*D*) alone, then of diameter and height (*H*) combined, by log-log regression (power law; Chave et al., 2014; Niklas, 1994) :

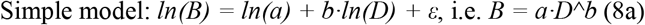

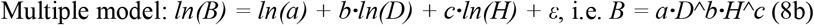

The two models were compared by *AIC* (Equation 6, applied here directly to the regression residuals on the logarithmic scale, both models sharing the same transformed response variable, which makes the comparison directly valid). The retained model is the one that minimises *AIC* while maximising adjusted *R*^*2*^ (Chave et al., 2014; Niklas, 1994).

All analyses were conducted in *R* and *Python 3*.*12* (pandas, numpy, scipy.optimize, statsmodels libraries), with a significance threshold set at α = 0.05 for all tests. The complete scripts (*Python* and *R*) are available and will be provided upon reasonable request to the corresponding author to ensure full reproducibility of the study.

## Results

### Germination

#### Overall phenology

Overall, germination of *Pterocarpus erinaceus* is epigeal, with the hypocotyl remaining in the soil; the epicotyl reaches an average length of 8.74 cm and an average basal diameter of 1.8 mm (on average 9 days after sowing). The prophylls (on average 10 days after sowing) have a long petiole oriented in the same direction as the hypocotyl, each bearing two (2) stipules at the base of the leaf sheath. The cotyledonary leaves remain visible until an average of 58 days. Phyllotaxis is alternate-spiral, with simple leaves bearing stipules until an average of 84 days, after which the seedling produces imparipinnate compound leaves, followed later by paripinnate compound leaves (on average 100 days). Branching appears in some seedlings after the emergence of compound leaves (on average 118 days).

#### A present but non-significant site effect

Cumulative germination kinetics differ little between sites (Figure 2). The final rate reached 70.3 ± 3.9% at Daloa (standard error, *n*=138 seeds) versus 61.6 ± 4.1% at Korhogo (*n*=138 seeds) (χ^*2*^=1.95; *p*=0.162), a difference that does not reach the significance threshold. Temporal dynamics are otherwise very similar between the two sites: germination reaches 50% of its final rate after *T50*=8.4 days at Daloa versus *T50*=8.6 days at Korhogo, a difference of only 0.2 days. This near-identical kinetics between sites, combined with the absence of a significant difference in final rate, suggests that germination in this species is relatively insensitive to the pedoclimatic contrast tested here, unlike its post-germination growth.

**Figure 2.**
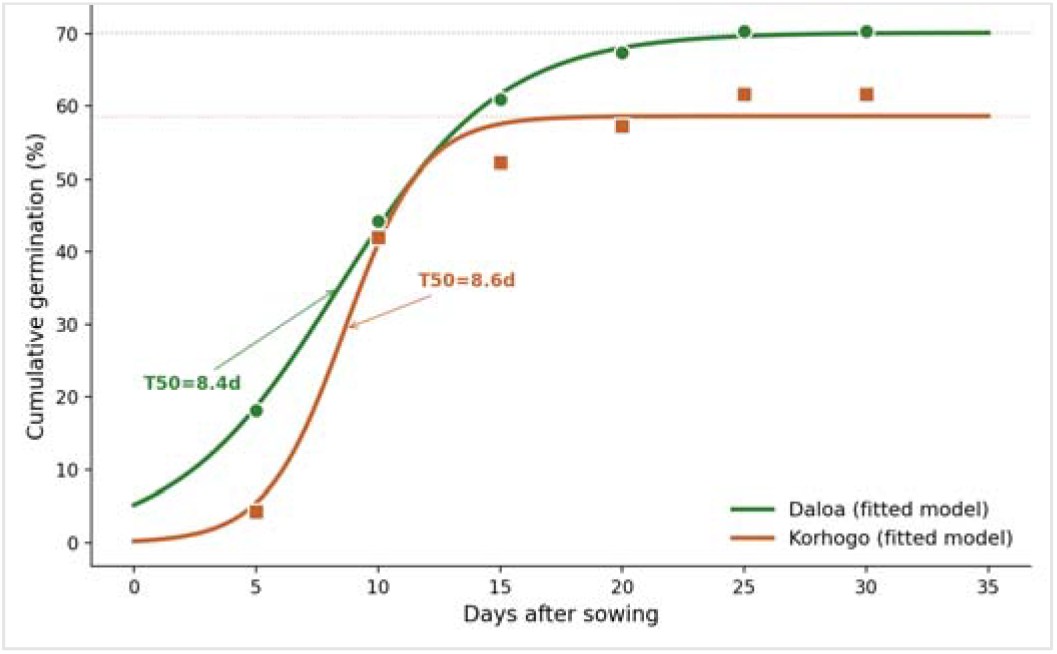
Cumulative germination kinetics of *Pterocarpus erinaceus* by study site, with fitted logistic models (*Gmax* = asymptotic germination rate; *T50* = time to reach 50% of final germination).

Analysis of variance (Table 2) reveals a significant site effect on all germination parameters (*P*<0.05), with the exception of germination spread duration, which is statistically equivalent between the two sites (*P*>0.05). Seed-tree group characteristics, by contrast, have no significant influence on seed germination, whatever the parameter considered (*P*>0.05 for all five parameters, sites combined). These germination parameters (Table 2), derived from the main six-seed-tree-group design, are independent of the cumulative kinetics presented in Figure 2, which relies on a distinct sub-sample used to characterise daily germination dynamics across all seed sizes combined.

**Table 2.** Overall influence of study sites and seed-tree groups on germination.

| Sites/Seed-tree group | Lag time | Germination delay | Germination rate | Germination duration | Germination percentage |
| --- | --- | --- | --- | --- | --- |
| Korhogo | $3.50 \pm 0.22$ b | $8.59 \pm 0.58$ b | $10.50 \pm 0.76$ b | $10.17 \pm 0.48$ a | $70.41 \pm 3.70$ b |
| Daloa | $3.83 \pm 0.40$ b | $7.16 \pm 0.29$ b | $6.83 \pm 0.40$ c | $8.17 \pm 0.47$ a | $87.22 \pm 2.34$ a |
| *Pr > F* | **0.0001** | **0.0004** | **0.001** | **0.1251** | **0.0001** |
| **Group-1** | 5.67 ± 2.18 **a** | 9.18 ± 2.91 **a** | 13.67 ± 6.23 **a** | 11.33 ± 2.33 **a** | 65.55 ± 12.37 **a** |
| **Group-2** | 5 ± 1 **a** | 8.19 ± 0.89 **a** | 10 ± 2.31 **a** | 7.67 ± 0.66 **a** | 66.66 ± 9.62 **a** |
| **Group-3** | 5.33 ± 1.45 **a** | 9.63 ± 1.23 **a** | 13.33 ± 4.84 **a** | 11 ± 2.08 **a** | 67.77 ± 10.59 **a** |
| **Group-4** | 6 ± 1.53 **a** | 9.47 ± 0.94 **a** | 12.33 ± 2.96 **a** | 8.67 ± 0.88 **a** | 71.99 ± 8.87 **a** |
| **Group-5** | 4.33 ± 1.33 **a** | 8.52 ± 0.74 **a** | 11 ± 2.65 **a** | 10 ± 0.58 **a** | 78.44 ± 10.05 **a** |
| **Group-6** | 4.33 ± 1.33 **a** | 9.57 ± 1.50 **a** | 12.33 ± 3.28 **a** | 10 ± 1.15 **a** | 77.50 ± 3.82 **a** |
| **Pr > F** | **0.9534** | **0.9762** | **0.9843** | **0.5011** | **0.8784** |

For each trait, values sharing the same letters are not statistically different at the 5% threshold.

### Growth: site outweighs seed-tree type

#### Morphology at four months of age

At 4 months of development, the results (Table 3) show that only height (*HtPl*) and internode length (*LgEntr*) differ significantly between sites (*P*<0.05), all other morphological parameters remaining statistically equivalent between sites (*P*>0.05). Regarding the effect of seed-tree group, sites combined, only leaf number (*NbrFe*) and leaf length (*LgFe*) differ significantly between seed-tree groups (*P*<0.05), the four other parameters (*HtPl, Dcol, LaFe, LgEntr*) remaining statistically equivalent between groups.

**Table 3.** Overall influence of study sites and seed-tree groups on seedling development.

| Sites/Seed-tree group | *HtPl* (cm) | *Dcol* (mm) | *NbreFe* | *LgFe* (cm) | *LaFe* (cm) | *LgEntr* (cm) |
| --- | --- | --- | --- | --- | --- | --- |
| **Korhogo** | 15.58±0.74 **a** | 2.50±0.11 **a** | 6.99±0.49 **a** | 5.04±0.31 **a** | 4.46±0.30 **a** | 3.15±0.22 **a** |
| **Daloa** | 12.11±0.86 **b** | 2.26±0.12 **a** | 6.39±0.40 **a** | 5.32±0.31 **a** | 4.99±0.36 **a** | 1.95±0.14 **b** |
| **Pr > F** | **0.0024** | **0.4292** | **0.4168** | **0.4455** | **0.5456** | **0.0014** |
| **Group-1** | 14.12±1.48 **a** | 2.33±0.31 **a** | 5.90±0.40 **b** | 5.27±0.34 **ab** | 5.17±0.59 **a** | 2.50±0.45 **a** |
| **Group-2** | 13.29±1.88 **a** | 2.08±0.31 **a** | 7.17±0.43 **ab** | 5.60±0.42 **a** | 5.14±0.49 **a** | 3.16±0.55 **a** |
| **Group-3** | 14.93±1.90 **a** | 2.45±0.19 **a** | 7.51±0.05 **ab** | 5.58±0.12 **a** | 5.15±0.34 **a** | 3.31±0.55 **a** |
| **Group-4** | 15.69±0.64 **a** | 2.42±0.10 **a** | 5.98±0.19 **b** | 4.91±0.39 **ab** | 4.09±0.49 **a** | 2.20±0.17 **a** |
| **Group-5** | 15.78±1.66 **a** | 2.65±0.18 **a** | 8.12±0.31 **a** | 4.92±0.35 **ab** | 4.68±0.44 **a** | 3.10±0.46 **a** |
| **Group-6** | 14.15±1.81 **a** | 2.69±0.09 **a** | 6.42±0.66 **b** | 4±0.13 **b** | 3.84±0.03 **a** | 3.22±1.02 **a** |
| **Pr > F** | **0.8629** | **0.4272** | **0.0094** | **0.0335** | **0.1900** | **0.7129** |

For each trait, values sharing the same letters are not statistically different at the 5% threshold. *HtPl*: Seedling height; *Dcol*: Seedling collar diameter; *NbreFe*: Number of leaves; *LgFe*: Leaf length; *LaFe*: Leaf width; *LgEntr*: Internode length; *cm*: centimetres; *mm*: millimetres.

#### Growth and modelling at 6, 12 and 24 months

##### Site, not seed-tree type, determines growth

Three independent and convergent statistical approaches demonstrate that the growth of *P. erinaceus* is determined by the planting site rather than by the dendrometric characteristics of the original seed tree.

#### Analysis of variance and boxplot

A classical ANOVA confirms the magnitude of the site effect on the three growth parameters (height, collar diameter and total dry biomass) at each age (Table 4; Figure 3). Biomass was not measured at 12 months for the reasons set out in the methodology (anomaly detected in this measurement file); height and diameter data at this age remain reliable and are reported in the table below.

**Table 4.** Mean ± standard deviation of the main growth parameters of *Pterocarpus erinaceus* by site and age.

| Age | Parameter | Daloa (mean±SD) | Korhogo (mean±SD) | *p* |
| --- | --- | --- | --- | --- |
| 6 months | Height (cm) | 19.8±5.3 | 13.3±2.8 | <0.001 |
| 6 months | Diameter (mm) | 4.56±0.78 | 3.32±1.03 | <0.001 |
| 6 months | Biomass (g) | 11.07±2.49 | 5.14±1.81 | <0.001 |
| 12 months | Height (cm) | 60.0±31.1 | 21.5±4.9 | <0.001 |
| 12 months | Diameter (mm) | 8.57±2.98 | 3.81±0.7 | <0.001 |
| 24 months | Height (cm) | 296.4±63.1 | 77.3±14.0 | <0.001 |
| 24 months | Diameter (mm) | 34.93±7.85 | 21.80±6.82 | <0.001 |
| 24 months | Biomass (g) | 498.3±238.0 | 67.3±19.2 | <0.001 |

**Figure 3.**
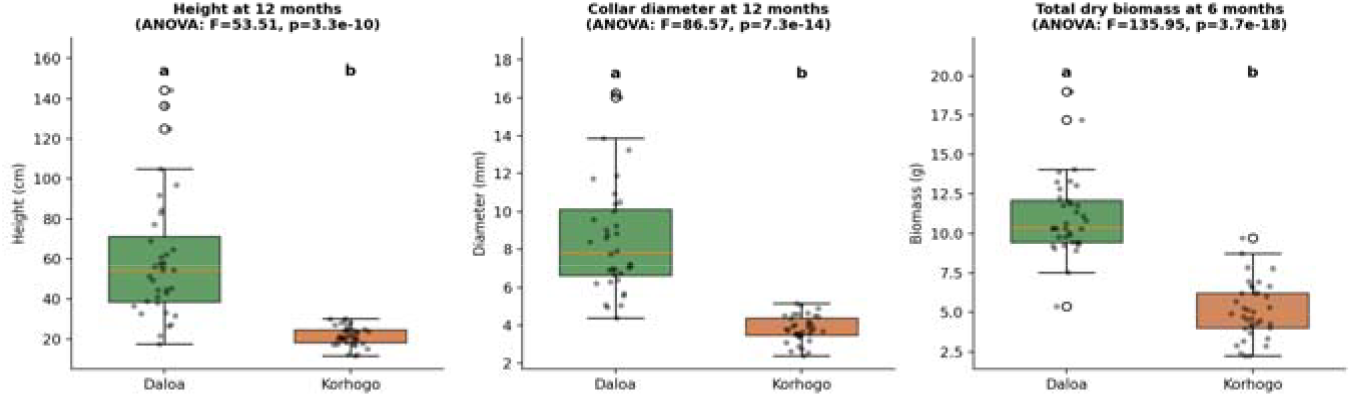
Boxplots showing the effect of study site on height, collar diameter, and total dry biomass of *Pterocarpus erinaceus*, with ANOVA results (letters denote significant differences, *p*<0.05).

For comparison, Figure 4 shows the effect of seed-tree group on height at 12 months: no significant difference appears among the six groups, in contrast to the substantial site effect observed above.

**Figure 4.**
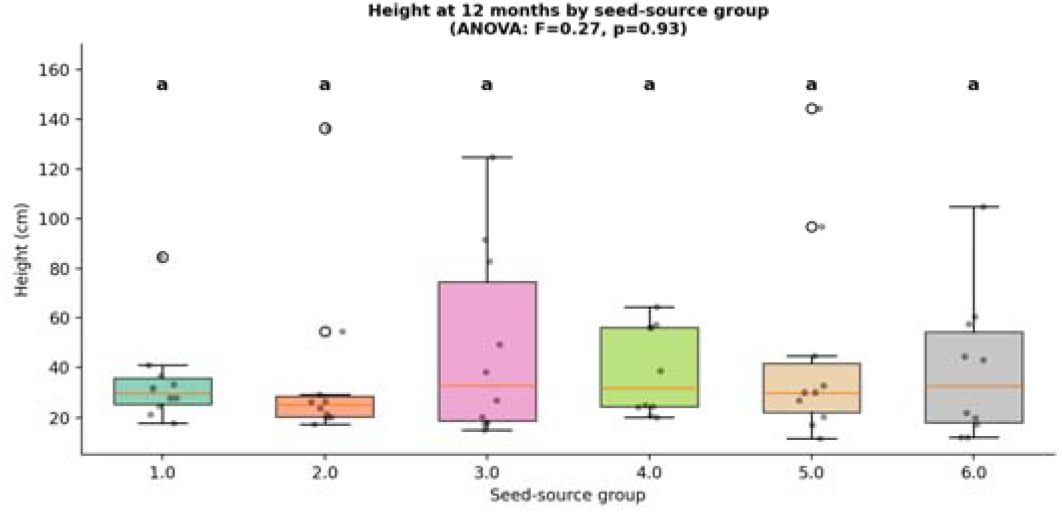
Height at 12 months by seed-source group: no significant difference between mother-tree groups (boxplot with ANOVA).

#### Continuous regression

Rather than comparing seed-tree categories, each growth parameter was regressed directly on the actual DBH and height of the original seed tree, separately at each age (Figure 5). Of the 16 regressions tested and performed (3 parameters × 3 ages × 2 predictors, excluding biomass at 12 months, excluded for the reasons set out in the methodology), 15 are not significant (*p* between 0.14 and 0.98). Only one reaches significance: collar diameter at 24 months is associated with seed-tree height (*r*^*2*^=0.22; *p*=0.015; *n*=26). With 16 tests conducted at the 5% threshold, approximately 0.8 (16 × 0.05) falsely significant result is statistically expected by chance alone. This isolated result should therefore be interpreted with caution rather than as proof of a robust biological effect, particularly as it is not found for either height or biomass at other ages.

**Figure 5.**
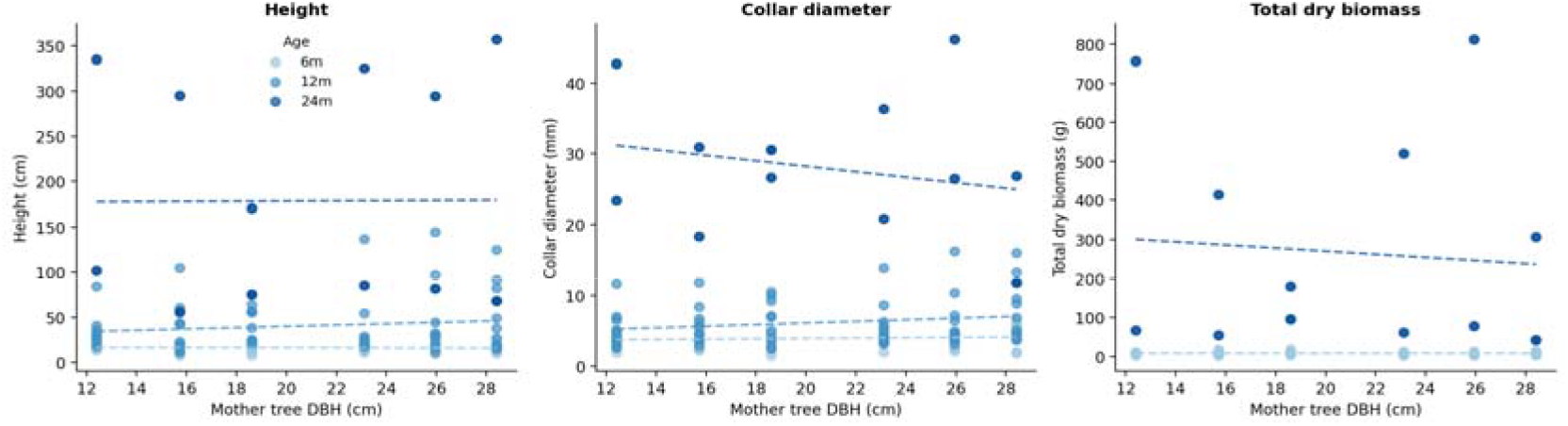
Continuous regression of growth parameters against mother tree DBH, by age class. Of the 16 regressions tested, only collar diameter at 24 months shows a significant association, with mother-tree height (*p*=0.015; *R*^*2*^=0.22; *n*=26); all other relationships are non-significant (*p*>0.14).

#### Linear mixed models with interaction: site × seed-tree group

Linear mixed models were fitted for height, collar diameter and total biomass (log-transformed), with seed tree as a random effect, site, age and seed-tree height as fixed effects, and a DBH × site interaction term, allowing a test of whether the seed-tree effect differs according to the environment (Table 5).

**Table 5.**
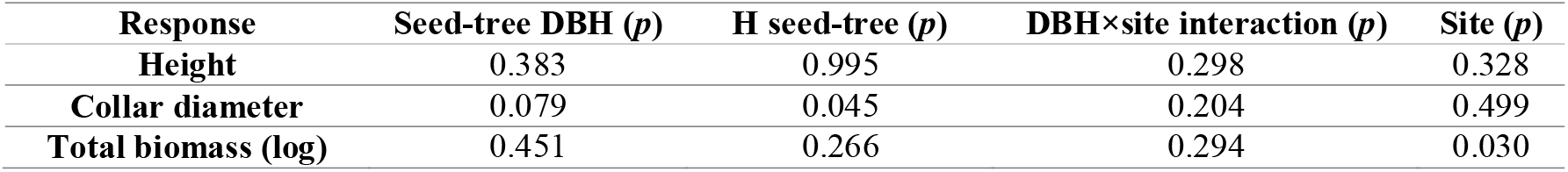
Results of linear mixed models (seed tree as random effect) for three growth responses.

Age is highly significant in both models (*p*<0.005 for the 12 months vs 6 months and 24 months vs 6 months comparisons), confirming the expected progression of growth over time; this effect, not central to the seed-tree question, is not detailed in Table 5. Seed-tree DBH is not significant in any of the three models. Seed-tree height, by contrast, shows a significant effect on collar diameter (*p*=0.045), consistent with the isolated result from the continuous regression. This signal, although reproduced by two independent methods, remains of small magnitude and confined to a single parameter-predictor combination out of the nine tested in this table.

#### Absence of an indirect effect via seed mass: mediation analysis

An alternative hypothesis is that seed-tree size could indirectly influence the performance of its offspring through the mass of the seeds it produces (a classical maternal effect in seed ecology). To determine whether seed-tree type acts via seed mass, a mediation pathway was tested at the level of the six seed-tree groups: seed-tree DBH → mean seed mass → germination rate → mean seedling height (Figure 6).

**Figure 6.**
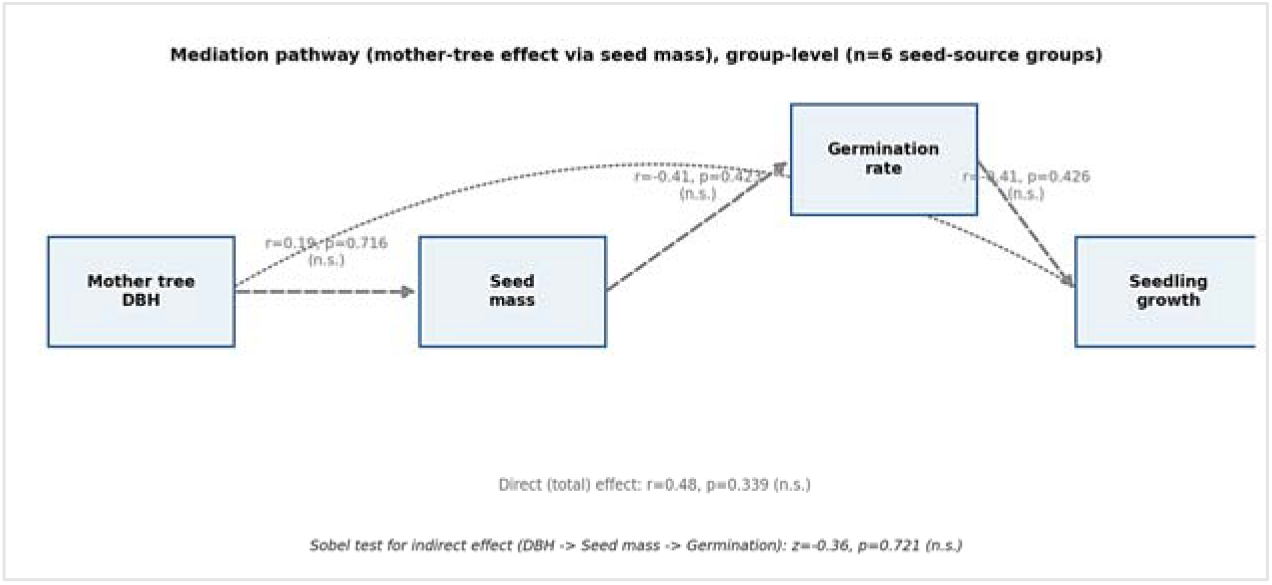
Mediation path linking mother tree DBH to offspring growth through seed mass and germination rate (group-level analysis, *n*=6 seed-source groups).

Seed mass in Pterocarpus erinaceus varies very little among seed trees within the same population (0.07 to 0.08 g, *Pr>F*=0.851 in the original analysis of variance), which strongly limits the power of this analysis: no link in the pathway is significant (DBH → seed mass: *r*=0.19, *p*=0.72; seed mass → germination: *r*=-0.41, *p*=0.42; germination → growth: *r*=-0.41, *p*=0.43), and the Sobel test for the indirect effect is not significant (*z*=-0.36; *p*=0.72). This lack of result should be interpreted with caution given the low variance in seed mass in this species, rather than as definitive proof of an absence of mediation.

#### Non-linear modelling of growth trajectories

Three growth models (linear, power law, exponential) were fitted to individual height and collar-diameter data, separately by site, and compared by *AIC* calculated on a common scale (Tables 6 and 7). A Gompertz model was also tested on the individual height data, but its asymptote parameter proved extremely poorly constrained (relative error of approximately 300%), the three available age classes (6-24 months) covering only an ascending growth phase, with no indication of a slowdown allowing a plateau to be estimated; this model was therefore not retained.

**Table 6.** Comparison of height growth models by *AIC* (common scale, see Figure 7).

| Site | Model | AIC | R<sup>2</sup> |
| --- | --- | --- | --- |
| Daloa | Linear | 623.2 | 0.830 |
| Daloa | Power law | 580.7 | 0.897 |
| Daloa | Exponential | 580.2 | 0.898 |
| Korhogo | Linear | 389.0 | 0.846 |
| Korhogo | Power law | 357.9 | 0.892 |
| Korhogo | Exponential | 331.3 | 0.920 |

**Table 7.** Comparison of collar-diameter growth models by *AIC* (common scale; see Figure 8).

| Site | Model | *AIC* | *R*<sup>2</sup> |
| --- | --- | --- | --- |
| Daloa | Linear | 259.9 | 0.815 |
| Daloa | Power law | 225.9 | 0.877 |
| Daloa | Exponential | 213.1 | 0.894 |
| Korhogo | Linear | 241.4 | 0.708 |
| Korhogo | Power law | 205.7 | 0.807 |
| Korhogo | Exponential | 189.7 | 0.839 |

##### Height growth

The exponential model is the best fit at Korhogo (Δ*AIC*=26.6 relative to the power law), but the advantage is minimal at Daloa (Δ*AIC*=0.5 only), where the two models are practically equivalent. The estimated parameters (Daloa: *H*=10.83×*exp*(0.138×*age*); Korhogo: *H*=6.57×*exp*(0.103×*age*)) reveal a significantly higher relative growth rate at Daloa (13.8%/month) than at Korhogo (10.3%/month), a relative difference of more than 30%. In *Pterocarpus erinaceus*, the growth gap in height between sites is therefore not explained solely by a difference in initial size, but also by an intrinsically faster growth pace at Daloa, suggesting that the edaphic and climatic conditions of this site favour faster stem elongation, and not merely better early seedling establishment (Table 6, Figure 7).

**Figure 7.**
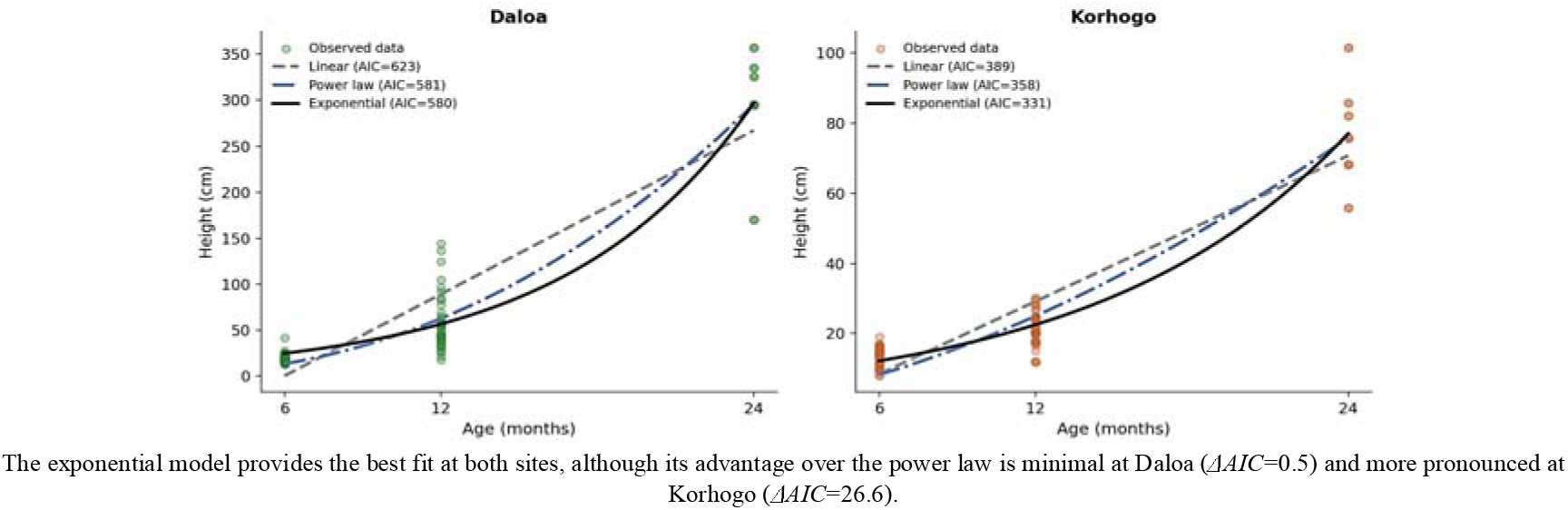
Comparison of the three growth models (linear, power law, exponential) fitted to individual height data of *Pterocarpus erinaceus*, by site.

##### Collar-diameter growth

The exponential model is also the best fit for collar diameter at both sites (Δ*AIC*=12.8 at Daloa and 16.0 at Korhogo relative to the power law). The estimated parameters (Daloa: *D*=2.18×*exp*(0.116×*age*); Korhogo: *D*=1.00×*exp*(0.128×*age*)) reveal a pattern opposite to that observed for height: the relative growth rate of diameter is slightly higher at Korhogo (12.8%/month) than at Daloa (11.6%/month), despite a markedly lower initial diameter at Korhogo. This divergence between the two organs suggests differential resource allocation depending on site: height growth, prioritised at Daloa, may come relatively at the expense of radial thickening, whereas at Korhogo, where conditions are more limiting for elongation, seedlings would invest proportionally more in diameter. This represents a strategy consistent with an allocation trade-off between vertical growth (light access) and structural stability (resistance to water stress and wind), already documented in other juvenile savanna woody species (Table 7, Figure 8).

**Figure 8.**
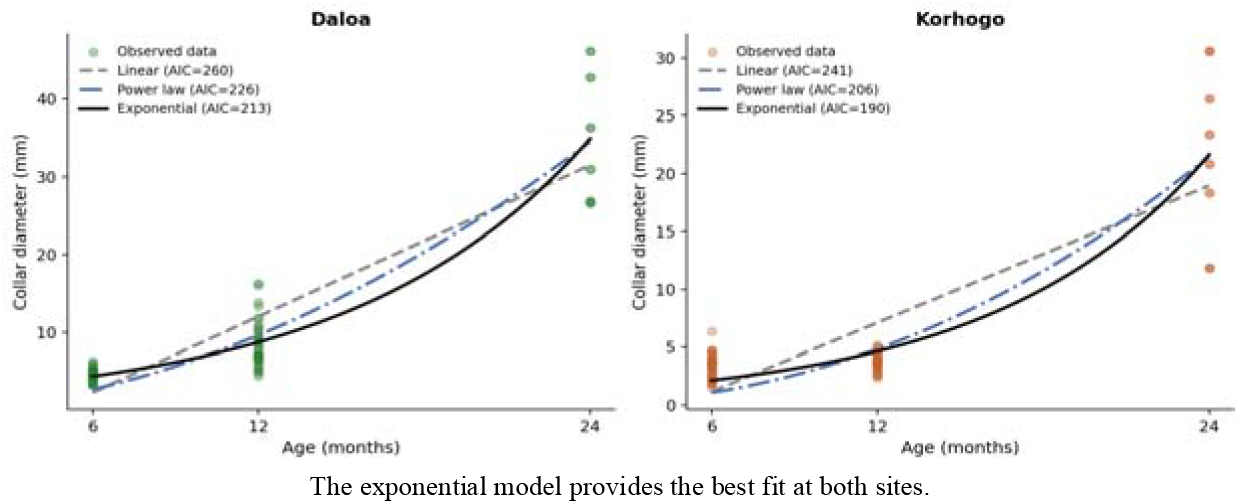
Comparison of the three growth models (linear, power law, exponential) fitted to individual collar-diameter data of *Pterocarpus erinaceus*, by site.

#### Between-site phenotypic plasticity

A between-site phenotypic plasticity index (RDPI, *Relative Distance Plasticity Index*) was calculated for height, by seed-tree group and age, based on the relative difference between the Daloa and Korhogo means (Figure 9). This index ranges from 0.05 to 0.68 depending on the group and age considered (a wider range than reported for other species in the same design), but this variation is correlated neither with seed-tree DBH (*r*=0.15; *p*=0.56, all data combined), nor significantly different among seed-tree groups (ANOVA: *F*=0.78; *p*=0.58). This result indicates that, while mean seedling performance varies markedly between sites, the extent of this variation (plasticity itself) likewise does not depend on the size of the original seed tree; a result that closes one last possible door to a hidden seed-tree effect. By contrast, this plasticity increases markedly with seedling age: the mean index rises from 0.19 ± 0.13 at 6 months to 0.46 ± 0.12 at 12 months, then to 0.57 ± 0.11 at 24 months, a progression strongly correlated with age (*r*=0.75; *p*<0.001; ANOVA: *F*=15.89; *p*<0.001). In other words, the performance gap between Daloa and Korhogo widens progressively over the course of juvenile development, rather than remaining stable from establishment onward. This represents a pattern consistent with a cumulative effect of the pedoclimatic contrast between the two sites, whose influence on growth intensifies month after month rather than manifesting immediately.

**Figure 9.**
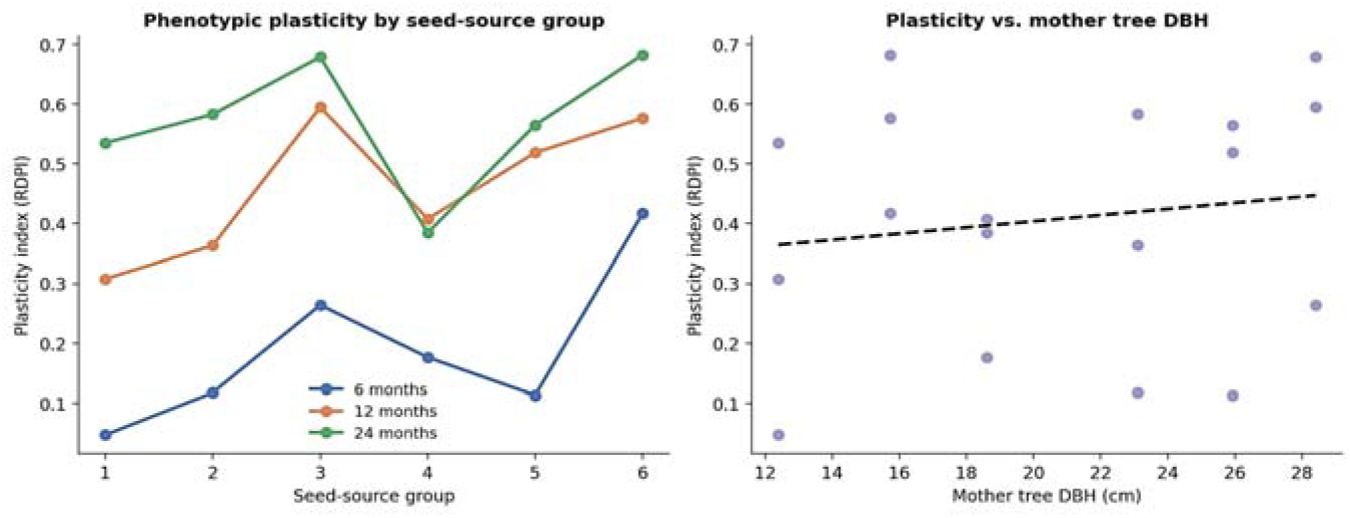
Between-site phenotypic plasticity index (RDPI) for height, by seed-source group and age, and its relationship with mother tree DBH.

#### Allometric equations and model comparison

The allometric relationship between collar diameter and total dry biomass is strong and highly significant (*R*^*2*^=0.90; Figure 10), following a power law with an exponent of 1.58; a more moderate exponent, reflecting biomass allocation less strongly concentrated on the diameter axis in this species. By contrast, the allometric equations for the leaf (*R*^*2*^=0.07) and root (*R*^*2*^=0.27) components remain weak, unlike the total biomass-diameter relationship. This weakness is largely explained by the reduced dataset available for these two organs following the exclusion of 12-month biomass measurements and the correction of physically impossible root-mass values, and these two equations should therefore be considered preliminary rather than reliable prediction tools for this species. Leaf area, conversely, is well predicted by leaf length (*R*^*2*^=0.84), offering a robust non-destructive indicator despite the limitations affecting leaf biomass itself. The root-to-shoot ratio could only be reliably calculated at 6 months (0.91 ± 0.81 on average, *n*=59), data at 12 and 24 months being insufficient or affected by the anomalies already noted. This high ratio at an early stage suggests already substantial root investment in young seedlings of this species, but cannot be tracked over time with the data currently available.

**Figure 10.**
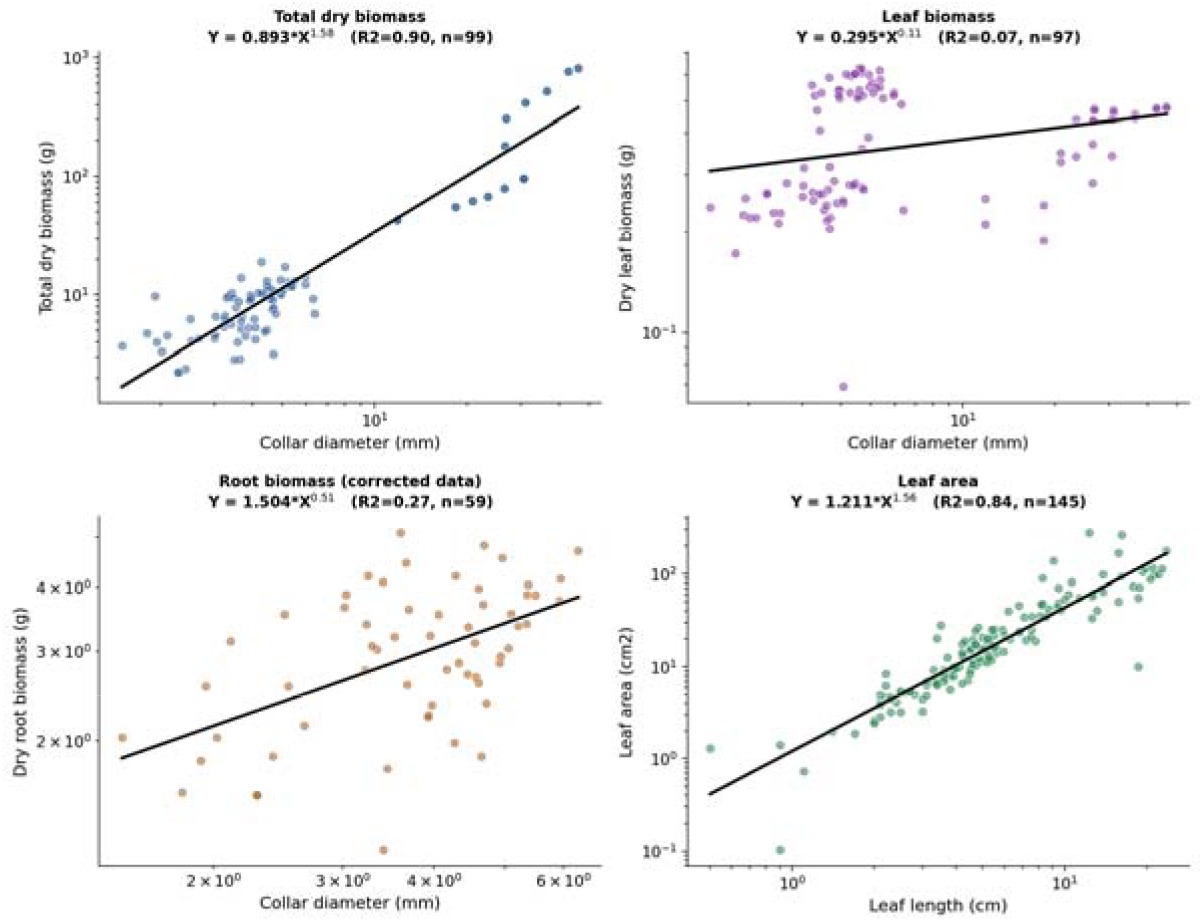
Allometric equations (log-log regression) relating collar diameter or leaf length to the various biomass components and to leaf area in *Pterocarpus erinaceus* seedlings (6-24 months, sites combined).

A single-predictor allometric model (total biomass ~ diameter alone) was then compared with a two-predictor model (diameter and height), by *AIC* calculated on a logarithmic scale (Table 8). The two-predictor model is markedly superior (Δ*AIC*=94.6 relative to the single-predictor model), with a substantial reduction in residual error (*R*^*2*^ raised to 0.962). The retained equation is: *ln(Biomass)* = −1.60 + 0.34×*ln(Diameter)* + 1.12×*ln(Height)*, providing a non-destructive estimation tool substantially more accurate than the single-predictor equation for nursery and reforestation-trial monitoring.

**Table 8.** Comparison of allometric models of total dry biomass, by *AIC* (logarithmic scale).

| Model | *AIC* | $R^2$ |
| --- | --- | --- |
| Power law (diameter only) | 141.4 | 0.899 |
| Quadratic log-log (diameter only) | 118.9 | 0.921 |
| Multiple allometry (diameter + height) | 46.8 | 0.962 |

#### Correlation structure of morphological traits

The correlation matrix (Figure 11) reveals strong phenotypic integration among aboveground growth traits: height, collar diameter, leaf area and total biomass are strongly correlated with one another (*r*=0.87 to 0.93). Root traits, by contrast, show markedly more moderate correlations with the other traits. Root length ranges from *r*=0.18 (with root biomass) to *r*=0.76 (with height), while root biomass itself is only weakly correlated with aboveground traits (*r*=0.21 to 0.50) and even shows a negative correlation with root length relative to total biomass (*r*=-0.35). This looser correlation structure for the root compartment is consistent with the reduction and correction of the dataset specifically affecting this organ, and calls for relationships involving root biomass to be interpreted with caution rather than as reflecting a genuine biological decoupling between aboveground and root growth in *Pterocarpus erinaceus*.

**Figure 11.**
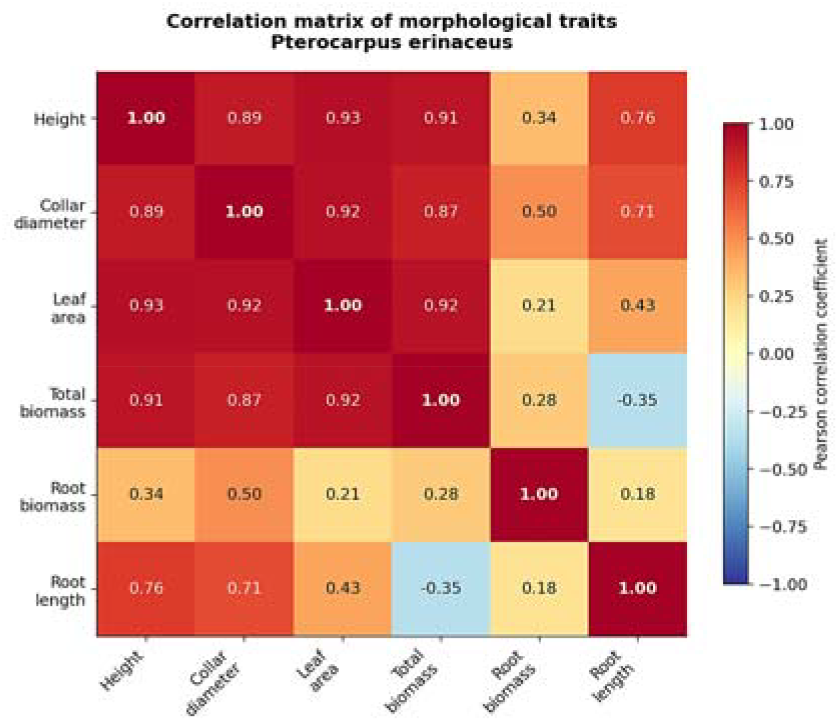
Pearson correlation matrix of the main morphological traits measured on *Pterocarpus erinaceus* seedlings.

## 4. Discussion

For an endangered timber species under severe harvesting pressure, should conservation programmes invest scarce resources in identifying large mother trees, or in choosing suitable planting environments? In the case of *Pterocarpus erinaceus* whose remaining mature trees are increasingly scarce, conservation programmes may gain more by matching planting material to suitable environments than by restricting seed collection to the largest surviving trees.

### Site, not seed tree, determines restoration success on Pterocarpus erinaceus

The results obtained here, converging across classical ANOVA, continuous regression and mixed models, confirm hypothesis H1 and largely corroborate hypothesis H2: the planting site very largely dominates the growth of *Pterocarpus erinaceus*, whereas no dendrometric characteristic of the seed tree has a detectable effect on height or biomass. This result directly extends that of Adji et al. (2021), who had already observed, using a design comparing three contrasting environments (Korhogo, Daloa, and a controlled greenhouse in Montpellier), a significant effect of environment on germination but no effect of the six sampled seed trees. A result obtained, however, through a simple categorical analysis of variance limited to the 4-month stage. The present study confirms this conclusion through continuous regression and mixed modelling, and extends it to 24 months: the convergence of these approaches, across two independent studies conducted by the same team a few years apart, constitutes a particularly robust body of evidence.

It is notable that these authors also reported a seed-tree effect limited to leaf number and length, with no effect on height, diameter or leaf width in 4-month-old seedlings. Our own results at this same stage (Table 3) confirm this pattern very precisely: only leaf number and leaf length differ significantly according to seed-tree group, while height, diameter, leaf width and internode length remain equivalent between groups. This near-exact replication, on a partially distinct sub-sample, reinforces the robustness of this exception localised to leaf dimensions, which may reflect an early plastic component more sensitive to small variations in germination conditions than structural traits (height, diameter) are, without calling into question the overall conclusion of an absence of effect on the offspring’s overall performance. This leaf-related exception is further part of a broader pattern: at 4 months, only two of six parameters (height, internode length) differed significantly between sites (Table 3), compared with the totality of parameters measured at 6, 12 and 24 months (Table 4). This still partial differentiation at 4 months, which becomes nearly total by 6 months, suggests that the site effect takes several months to fully manifest across all morphological traits, rather than acting from the very first weeks of growth. This result further corroborates the findings of Tiika et al. (2019) in Ghana, who found no correlation between seed mass and germination percentage according to mother-tree size in this same species. The convergence of these independent studies strongly reinforces the generality of the message: in *Pterocarpus erinaceus*, selecting seed trees on size criteria provides no measurable benefit for the regeneration and restoration of the species.

Germination, by contrast, shows no significant difference between sites in this species (*p*=0.162). This result echoes the work of Ansah et al. (2023), who showed that the characteristics of Pterocarpus erinaceus seeds (potentially their germination potential) respond more strongly to maternal environmental conditions (temperature, soil nutrients at the seed-production site) than to the conditions of the nursery sowing site itself. It is also possible that the edaphic variability between Daloa and Korhogo, although sufficient to strongly affect post-germination growth, does not reach the threshold needed to modulate the germination process itself in this species, whose seeds are protected by a seed coat inducing physical dormancy relatively independent of the substrate. This observation is consistent with the findings of Asigbaase et al. (2024) in Ghana, who showed that the survival and growth of *P. erinaceus* vary strongly between contrasting ecological zones, without an equivalent effect being consistently reported for germination. Analysis of variance applied to the detailed germination parameters (Table 2) refines this picture: site significantly influences lag time, delay and germination rate (*P*<0.05), but not the spread of the germination process (*P*>0.05), while none of these five parameters differs according to seed-tree group. This dissociation (a selective site effect on certain kinetic parameters but not on the overall temporal spread of germination) suggests that site mainly modulates the speed of initiation of the germination process rather than its total duration, consistent with the absence of a difference in final rate already noted for the pooled kinetics.

One isolated signal nonetheless warrants cautious discussion: seed-tree height is associated with collar diameter at 24 months, both in simple regression and in the mixed model. Given the number of tests performed (16 regressions, 9 mixed-model coefficients), this single result could reflect a statistical false positive rather than a genuine biological effect; it is not reproduced for either height or biomass, nor at other ages. We report it in the interest of full scientific transparency, but recommend its replication in an independent design before any causal interpretation.

### A near-invariant seed mass: a species-specific biological constraint

The mediation analysis, although inconclusive, highlights an important point: in *Pterocarpus erinaceus*, seed mass varies almost not at all among seed trees within the same population (0.07 to 0.08 g, *Pr>F*=0.851 in the original analysis of variance). This structural near-invariance limits any possibility of detecting a maternal effect through this pathway in this species, consistent with hypothesis H3. This finding underscores that the absence of mediation observed here likely reflects a species-specific biological constraint (perhaps linked to an indehiscent pod imposing a relatively fixed seed size), rather than a conclusion generalisable to other woody Fabaceae.

### Rapid juvenile growth, contrasting with an extremely slow adult regime

The exponential model consistently dominates for both height and diameter, at both sites, with relative growth rates on the order of 10 to 14% per month depending on site and organ. This rapid juvenile growth pace contrasts strikingly with the estimates available for the adult phase of the species: the IUCN Red List assessment (Barstow, 2017) uses a height growth rate of approximately 15 cm per year in adulthood (barely more than what our seedlings gain in a single month at Daloa); the species requiring nearly a century to reach its adult height of 15 m. This discontinuity between a rapid juvenile phase and an extremely slowed adult regime underscores the critical importance of the first two years following germination for any active restoration intervention: it is during this window, when growth potential is at its maximum, that the choice of site produces most of its effect, before the species enters a much slower growth regime in which differences between individuals or provenances may become harder to correct.

The differential allocation observed between height and diameter according to site (height growth relatively favoured at Daloa, diameter at Korhogo) is consistent with a classical allocation trade-off between light access and structural stability, already documented in tropical tree seedlings facing contrasting water and light constraints (Markesteijn and Poorter, 2009). Furthermore, the homogeneous exponential model for both height and diameter at both sites suggests that *Pterocarpus erinaceus* has not yet, by 24 months, entered a phase of radial growth deceleration, consistent with an architectural development strategy in which investment in height and diameter remains coupled over this juvenile developmental window.

### Implications for conservation and study limitations

This study supports prioritising the choice of planting site over the dendrometric selection of seed trees in restoration programmes for *Pterocarpus erinaceus*, a species whose conservation urgency is heightened by its CITES listing and the international commercial pressure it faces (Dumenu, 2019). These results have direct practical relevance for ex situ conservation and compensatory reforestation programmes mandated in some exporting countries (CITES, 2018) in response to the rosewood trade crisis. The population decline estimated at 80% in the Gambia, Benin and Côte d’Ivoire (Barstow, 2017; cfeditorfr, 2022; IUCN, 2018) illustrates the urgency of these interventions and the growing scarcity of mature seed trees available for seed collection. In this context of limited genealogical resources, our results are reassuring on one specific point: managers do not need to prioritise the few surviving large seed trees at the expense of the accessibility or genetic diversity of the material collected. Priority should instead go to the choice of planting site, a factor over which practitioners have direct and immediate control.

This study has several limitations. The data anomaly detected at 12 months (74 out of 145 individuals with root data, i.e. 51%) calls for a thorough review of the biomass measurement protocol used during that campaign (possibly linked to a balance calibration issue or a systematic transcription error). This limitation mainly affects the precision of the root and leaf allometric equations, which rely on a reduced sample after exclusion of suspect data. The number of seed-tree groups (six) inherently limits the power of the mediation analysis. The monitoring window (24 months) does not allow characterisation of the transition to the slowed growth regime expected in adulthood, and the Gompertz model could not be reliably fitted owing to the lack of observations covering this deceleration phase. An exploratory test, incorporating an additional anchor point at 7 years (*n*=12 individuals, Daloa site only) into the 6-24-month height data, suggests that a Gompertz model could technically be fitted over this extended window, with an apparently more favourable *AIC* than the two-parameter models. Finally, this study was conducted at only two sites; replication across a larger number of sites would allow the conclusion to be generalised to the species’ entire Sudano-Guinean distribution range.

## Conclusion

While barwood continues to disappear at an alarming rate, with populations having declined by nearly 80% under the pressure of illegal international rosewood trade, this study provides a clear answer to a question directly actionable for compensatory propagation and ex situ conservation programmes: it is not seed-tree size, but the choice of planting site, that constitutes the major determinant of juvenile growth in *Pterocarpus erinaceus*. Converging across the full range of analytical approaches used, from site comparison to formal mediation, this result methodically closes off the avenues through which a seed-tree effect might have manifested, with the exception of an isolated and statistically fragile signal between seed-tree height and diameter, to be interpreted with caution. In a context of accelerating scarcity of mature seed trees across the species’ range, managers can be reassured: strict selection based on size criteria is not necessary to produce vigorous seedlings. Priority should instead go to the choice of planting site. The striking contrast between the rapid juvenile growth observed here and the extremely slow adult growth regime reported in the literature further underscores that the first two years of life constitute a critical window for any active conservation intervention. The multi-predictor allometric equations developed here also provide a non-destructive monitoring tool directly applicable in conservation nurseries. Ultimately, this study calls for refocusing barwood restoration programmes on a simpler, yet more decisive, question than seed-tree selection: *where to plant, rather than which seeds to choose*.

## Supporting information

Supplementary_Scripts python and R for Pterocarpus

## CRediT authorship contribution statement

**Beda Innocent Adji**: Conceptualization, Data curation, Formal analysis, Investigation, Methodology, Resources, Software, Validation, Visualization, Writing – original draft, Writing - review & editing.

## Fundind

The work described in this document was carried out using the author’s own funds. No funding was received from any funding body for the supervision and execution of the fieldwork (tree sampling, seed collection by locality, and nursery trials and monitoring).

## Declaration of competing interest

The author declare that they have no known competing financial interests or personal relationships that could have appeared to influence the work reported in this paper.

## Acknowledgements

The author would like to thank the rural people of the various localities (Katiola, Niakara, Korhogo, and Sinématiali) visited for their mobilisation, generosity, and hospitality during the phases of seed provenance collection and mother-tree sampling. The author thank the various owners of the agroforestry parklands visited for making their orchards available and granting permission to conduct this study on the various trees within them. Finally, the author thank the *Centre National de Recherche Agronomique* of Côte d’Ivoire (CNRA) for making available the Diabaté Kamonon forestry research station in Sédiakaha, Korhogo, for part of this study.

## Data availability

The datasets analysed as part of this study are available at: ADJI, B. I. (2026). Supporting Dataset for the document of Germination, morphology and biomass of *Khaya senegalensis, Pterocarpus erinaceus* and *Parkia biglobosa* seedlings according to mother-tree dendrometric characteristics and planting site in Côte d’Ivoire (2018-2025). https://doi.org/10.5281/zenodo.21435509

## Notes

https://doi.org/10.5281/zenodo.21435509

### Competing Interest Statement

The authors have declared no competing interest.

