## Supplementary_Scripts python and R for Pterocarpus for "Facing West Africa’s rosewood crisis: what determines the success of preserving and propagating the endangered barwood (*Pterocarpus erinaceus*)? A multi-model test of site and maternal-origin effects"

**Supplementary Information**

S1. Python script (produced all reported results)

The script below was executed in full to generate every statistical result, table value, and figure reported in the main text (data pipeline, quality control, germination kinetics, classical ANOVA, continuous regression, linear mixed models, mediation analysis with Sobel test, non-linear growth models including the tested Gompertz model and the exploratory 7-year extension, phenotypic plasticity index, allometric equations, and the correlation matrix).

```
# ==========================================================================
# Pterocarpus erinaceus - Complete analysis pipeline (Python 3.12)
# Site vs seed-tree effects on germination and juvenile growth
# ==========================================================================
# Libraries: pandas, numpy, scipy, statsmodels, openpyxl
# Raw data files: Pterocarpus_6_mois__morpho-biomasse_.xlsx
#               Pterocarpus_12_mois__morpho-Biomasse_.xlsx
#               Pterocarpus_24_mois__morpho-biomasse_.xlsx
#
GERMINATION_ET_DEVELOPPEMENT_SELON_taille_de_graine_chez_Khaya__Pterocarpus_et_Parkia.xlsx
#               Donnees_semenciers_7ans.xlsx (exploratory Gompertz extension only)
# ==========================================================================

import pandas as pd, numpy as np
import openpyxl, re
import statsmodels.api as sm
import statsmodels.formula.api as smf
from scipy import stats
from scipy.optimize import curve_fit
import warnings
warnings.filterwarnings('ignore')

# ----------------------------------------------------------------------
# 1. DATA PIPELINE: loading and reshaping raw growth files
# ----------------------------------------------------------------------

def load_sheet(path, sheet, header_row_idx):
    """Load one Excel sheet, using row header_row_idx (0-indexed) as column names."""
    wb = openpyxl.load_workbook(path, data_only=True)
    ws = wb[sheet]
    rows = list(ws.iter_rows(values_only=True))
    header = rows[header_row_idx]
    data = rows[header_row_idx + 1:]
    cols, seen = [], {}
    for i, h in enumerate(header):
        name = str(h).strip() if h else f"col{i}"
        if name in seen:
            seen[name] += 1
            name = f"{name}__dup{seen[name]}"
        else:
            seen[name] = 0
        cols.append(name)
    df = pd.DataFrame(data, columns=cols)
    df = df.loc[:, ~df.columns.str.startswith('col')]
    df = df.loc[:, ~df.columns.str.contains('__dup')]
    df = df.rename(columns={'semencier': 'semenciers'})
    header_like = df.apply(
        lambda r: any(str(v).strip().lower() in
                    ('semenciers', 'individus', 'semencier', 'localite', 'localites')
                    for v in r.values if pd.notna(v)), axis=1)
    df = df.loc[~header_like].reset_index(drop=True)
```

```
    return df

PLANT_LEVEL_CANDIDATES = [
    'long plant', 'diam plant', 'nbr UC', 'nbr module', 'nbr entr', 'long ptiol',
    'diam ptiol', 'long rachis', 'nbr fliol', 'nbr fliolul', 'long feuil',
    'larg feuil', 'long +grd fliol', 'larg +grd fliol', 'log +grd fliolul',
    'lag +grd fliolul', 'surface feuil', 'long raci', 'diam raci',
    'mas fr to plant', 'mas sc to plant', 'pd fr entr', 'pd sc entr',
    'pd fr feuil', 'pd sc feuil', 'LDMC', 'Pd fr ptiol', 'Pd sc ptiol',
    'pd fr fliol', 'pd sc fliol', 'pd fr fliolul', 'pd sc fliolul',
    'mas surfacique', 'pd fr raci', 'pd sc raci',
]
VARYING_CANDIDATES = ['long entr', 'diam entr']

def agg_individual(df, site, age):
    """Collapse per-organ measurement rows into one row per individual seedling."""
    df = df.copy()
    if 'individus' not in df.columns:
        key_cols = [c for c in ['long plant', 'diam plant', 'mas fr to plant'] if c in df.columns]
        change = (df[key_cols] != df[key_cols].shift()).any(axis=1)
        df['individus'] = df['semenciers'].astype(str) + '-ind' + change.cumsum().astype(str)
    agg_dict = {}
    for c in PLANT_LEVEL_CANDIDATES:
        if c in df.columns:
            agg_dict[c] = 'first'
    for c in VARYING_CANDIDATES:
        if c in df.columns:
            df[c] = pd.to_numeric(df[c], errors='coerce')
            agg_dict[c] = 'mean'
    g = df.groupby(['semenciers', 'individus'], dropna=False).agg(agg_dict).reset_index()
    g['localite'] = site
    g['age_mois'] = age
    return g

def extract_group(x):
    m = re.search(r'S-?(\d)', str(x))
    return int(m.group(1)) if m else None

def build_master(files, mother_tree_df):
    all_parts = []
    for age, (path, sD, sK, hdr) in files.items():
        dD = load_sheet(path, sD, hdr)
        dK = load_sheet(path, sK, hdr)
        all_parts.append(agg_individual(dD, 'Daloa', age))
        all_parts.append(agg_individual(dK, 'Korhogo', age))
    master = pd.concat(all_parts, ignore_index=True)
    master['groupe_semencier'] = master['semenciers'].apply(extract_group)
    master = master.merge(mother_tree_df, on='groupe_semencier', how='left')
    for c in master.columns:
        if c not in ['semenciers', 'individus', 'localite']:
            master[c] = pd.to_numeric(master[c], errors='coerce')
    return master

# Mother-tree dendrometric characteristics (Table 1)
mother_tree = pd.DataFrame({
    'groupe_semencier': [1, 2, 3, 4, 5, 6],
    'DBH_semencier_cm': [12.4, 23.11, 28.41, 18.6, 25.94, 15.72],
    'H_semencier_m':    [8.31, 14.7, 17.65, 12.23, 13.59, 15.66],
    'Age_semencier_ans':[23, 26, 26, 22, 25, 25],
    'Masse_graine_moy_g':[0.07, 0.08, 0.07, 0.07, 0.07, 0.07],
})

files = {
```

```
    6:  ('Pterocarpus_6_mois__morpho-biomasse_.xlsx',  'Daloa', 'Korhogo', 1),
    12: ('Pterocarpus_12_mois__morpho-Biomasse_.xlsx', 'Daloa', 'Korhogo', 1),
    24: ('Pterocarpus_24_mois__morpho-biomasse_.xlsx', 'Morpho-biom-Daloa', 'Morpho-biom-Korhogo', 1),
}
master = build_master(files, mother_tree)

# -------------------------------------------------------------------
# 2. DATA QUALITY CONTROL (section 2.6)
# -------------------------------------------------------------------

def audit_and_clean(master, biomass_col='mas sc to plant', root_col='pd sc raci',
                    root_fr_col='pd fr raci', threshold=0.8):
    """Flag individuals whose root dry mass exceeds total dry mass (physically impossible)."""
    sub = master.dropna(subset=[root_col, biomass_col]).copy()
    sub = sub[sub[biomass_col] > 0]
    ratio = sub[root_col] / sub[biomass_col]
    bad_idx = sub[ratio > threshold].index
    print(f"Quality control: {len(bad_idx)} root-mass values set to NaN (ratio>{threshold*100:.0f}%)")
    master.loc[bad_idx, root_col] = np.nan
    if root_fr_col in master.columns:
        master.loc[bad_idx, root_fr_col] = np.nan
    return master, len(bad_idx)

master, n_corrected = audit_and_clean(master)

# 12-month biomass anomaly: mean biomass at 12 months < mean at 6 months (impossible)
# -> all biomass-related columns at 12 months are excluded from analysis
biomass_cols = [c for c in master.columns if ('mas' in c.lower() or 'pd ' in c.lower()
                or 'LDMC' in c) and c != 'Masse_graine_moy_g']
mask_12mo = master.age_mois == 12
master.loc[mask_12mo, biomass_cols] = np.nan

master.to_csv('pterocarpus_master_clean.csv', index=False)

# -------------------------------------------------------------------
# 3. GERMINATION KINETICS (pooled cumulative curve, Figure 2)
# -------------------------------------------------------------------

def logistic(t, Gmax, k, t50):
    return Gmax / (1 + np.exp(-k * (t - t50)))

days = np.array([5, 10, 15, 20, 25, 30], dtype=float)
# daily counts pooled across the three seed-size classes (from the seed-size germination file)
daloa_daily   = np.array([22+1+2, 15+10+11, 5+15+3, 4+4+1, 0+2+2, 0+0+0])
korhogo_daily = np.array([4+2+0, 27+20+5, 3+5+6, 2+2+3, 2+2+2, 0+0+0])
n_total = 138
cum_daloa = np.cumsum(daloa_daily)
cum_korhogo = np.cumsum(korhogo_daily)

table = np.array([[cum_daloa[-1], n_total - cum_daloa[-1]],
                  [cum_korhogo[-1], n_total - cum_korhogo[-1]]])
chi2, pval, dof, exp = stats.chi2_contingency(table)
print(f"Germination site effect: chi2={chi2:.2f}, p={pval:.4f}")
print(f"Daloa final rate: {cum_daloa[-1]/n_total*100:.1f}%  |  Korhogo: {cum_korhogo[-1]/n_total*100:.1f}%")

pct_daloa = cum_daloa / n_total * 100
pct_korhogo = cum_korhogo / n_total * 100
popt_d, _ = curve_fit(logistic, days, pct_daloa, p0=[70, 0.3, 12], maxfev=10000)
popt_k, _ = curve_fit(logistic, days, pct_korhogo, p0=[70, 0.3, 12], maxfev=10000)
print(f"Daloa Gmax={popt_d[0]:.1f}%, T50={popt_d[2]:.1f} days")
print(f"Korhogo Gmax={popt_k[0]:.1f}%, T50={popt_k[2]:.1f} days")

# -------------------------------------------------------------------
```

```
# 4. CLASSICAL ANOVA (site and age effects, Table 4)
# ---------------------------------------------------------------------

df = pd.read_csv('pterocarpus_master_clean.csv')

for age in [6, 12, 24]:
    sub = df[df.age_mois == age]
    for col, label in [('long plant', 'Height'), ('diam plant', 'Diameter'), ('mas sc to plant', 'Biomass')]:
        s = sub.dropna(subset=[col, 'localite'])
        groups = [s[s.localite == site][col].values for site in ['Daloa', 'Korhogo']]
        if all(len(g) > 1 for g in groups):
            f, p_val = stats.f_oneway(*groups)
            print(f"{age}m {label}: F={f:.2f} p={p_val:.2e}")

# ---------------------------------------------------------------------
# 5. AXIS 1 - CONTINUOUS REGRESSION (16 tests, Figure 5)
# ---------------------------------------------------------------------

results = []
for age in [6, 12, 24]:
    sub_age = df[df.age_mois == age]
    for param in ['long plant', 'diam plant', 'mas sc to plant']:
        for pred in ['DBH_semencier_cm', 'H_semencier_m']:
            s = sub_age.dropna(subset=[param, pred])
            if len(s) < 5:
                continue
            X = sm.add_constant(s[pred])
            m = sm.OLS(s[param], X).fit()
            results.append({'age': age, 'param': param, 'pred': pred,
                            'p': m.pvalues.iloc[1], 'R2': m.rsquared, 'n': len(s)})
res_df = pd.DataFrame(results)
print(res_df.round(4).to_string(index=False))
print(f"Number of regressions performed: {len(res_df)}")
print(f"Significant (p<0.05): {(res_df.p < 0.05).sum()}")

# ---------------------------------------------------------------------
# 6. AXIS 2 - LINEAR MIXED MODELS (Table 5)
# ---------------------------------------------------------------------
# NOTE: fitted here with statsmodels (Python), NOT lme4 (R) as stated in the
# manuscript text. See accompanying R script for the lme4/lmerTest equivalent.

for col, label in [('long plant', 'Height'), ('diam plant', 'Diameter')]:
    sub = df.dropna(subset=[col, 'DBH_semencier_cm', 'H_semencier_m', 'localite', 'age_mois', 'groupe_semencier'])
    md = smf.mixedlm(f'Q("{col}") ~ DBH_semencier_cm * localite + H_semencier_m + C(age_mois)',
                     sub, groups=sub['groupe_semencier'])
    mdf = md.fit()
    print(f"--- {label} ---")
    print(mdf.pvalues.round(4))

sub = df.dropna(subset=['mas sc to plant', 'DBH_semencier_cm', 'H_semencier_m', 'localite', 'age_mois',
'groupe_semencier'])
sub = sub[sub['mas sc to plant'] > 0].copy()
sub['log_biomass'] = np.log(sub['mas sc to plant'])
md = smf.mixedlm('log_biomass ~ DBH_semencier_cm * localite + H_semencier_m + C(age_mois)',
                 sub, groups=sub['groupe_semencier'])
mdf = md.fit()
print("--- Biomass (log) ---")
print(mdf.pvalues.round(4))

# ---------------------------------------------------------------------
# 7. AXIS 3 - MEDIATION ANALYSIS AND SOBEL TEST (Figure 6)
# ---------------------------------------------------------------------
```

```
germ_groups = pd.DataFrame({
    'groupe_semencier': [1, 2, 3, 4, 5, 6],
    'taux_germination': [65.55, 66.66, 67.77, 71.99, 78.44, 77.50],
})
growth_by_group = df.groupby('groupe_semencier')['long plant'].mean().reset_index() \
                    .rename(columns={'long plant': 'mean_height'})
path = mother_tree.merge(germ_groups, on='groupe_semencier').merge(growth_by_group, on='groupe_semencier')

def reg(x, y, label):
    X = sm.add_constant(x)
    m = sm.OLS(y, X).fit()
    b = m.params.iloc[1]; se = m.bse.iloc[1]; p_val = m.pvalues.iloc[1]
    r = np.corrcoef(x, y)[0, 1]
    print(f"{label}: r={r:.3f} p={p_val:.3f}")
    return b, se, p_val, r

a_b, a_se, a_p, a_r = reg(path['DBH_semencier_cm'], path['Masse_graine_moy_g'], "DBH->SeedMass")
b_b, b_se, b_p, b_r = reg(path['Masse_graine_moy_g'], path['taux_germination'], "SeedMass->Germ")
c_b, c_se, c_p, c_r = reg(path['taux_germination'], path['mean_height'], "Germ->Growth")
d_b, d_se, d_p, d_r = reg(path['DBH_semencier_cm'], path['mean_height'], "DBH->Growth(total)")

indirect = a_b * b_b
se_i = np.sqrt(b_b**2 * a_se**2 + a_b**2 * b_se**2)
z = indirect / se_i
p_sobel = 2 * (1 - stats.norm.cdf(abs(z)))
print(f"Sobel test: z={z:.2f} p={p_sobel:.3f}")

# ----------------------------------------------------------------------
# 8. AXIS 4 - NON-LINEAR GROWTH MODELS (Tables 6-7, Figures 7-8)
# ----------------------------------------------------------------------

def lin_f(t, a, b): return a + b * t
def pow_f(t, a, b): return a * t**b
def exp_f(t, a, b): return a * np.exp(b * t)
def gompertz(t, A, B, k): return A * np.exp(-B * np.exp(-k * t))
def aic_from_rss(rss, n, k): return n * np.log(rss / n) + 2 * k

for varname, col in [('Height', 'long plant'), ('Diameter', 'diam plant')]:
    print(f"=== {varname} growth models ===")
    for site in ['Daloa', 'Korhogo']:
        sub = df[df.localite == site].dropna(subset=[col, 'age_mois'])
        t = sub['age_mois'].values.astype(float)
        y = sub[col].values.astype(float)
        n = len(t)
        print(f"--- {site} (n={n}) ---")
        p0map = {'linear': [0, 10], 'power': [3, 1.3], 'exponential': [10, 0.1]} if col == 'long plant' \
              else {'linear': [0, 1], 'power': [1, 1], 'exponential': [2, 0.1]}
        for name, f in [('linear', lin_f), ('power', pow_f), ('exponential', exp_f)]:
            popt, _ = curve_fit(f, t, y, p0=p0map[name], maxfev=20000)
            rss = np.sum((y - f(t, *popt))**2)
            aic = aic_from_rss(rss, n, 2)
            r2 = 1 - rss / np.sum((y - y.mean())**2)
            print(f"  {name}: AIC={aic:.1f} R2={r2:.3f} params={np.round(popt,4)}")

# Gompertz model tested on individual height data (found poorly constrained, not retained)
for site in ['Daloa', 'Korhogo']:
    sub = df[df.localite == site].dropna(subset=['long plant', 'age_mois'])
    t = sub['age_mois'].values.astype(float)
    y = sub['long plant'].values.astype(float)
    try:
        p0 = [y.max() * 2, 5, 0.05]
        bounds = ([y.max(), 0.1, 0.001], [y.max() * 20, 100, 5])
        popt, pcov = curve_fit(gompertz, t, y, p0=p0, bounds=bounds, maxfev=50000)
```

```
        perr = np.sqrt(np.diag(pcov))
        print(f"{site} Gompertz: A={popt[0]:.1f}+-{perr[0]:.1f}, rel.err={perr[0]/popt[0]*100:.0f}%")
    except Exception as e:
        print(f"{site} Gompertz fit failed: {e}")

# --- Exploratory extension: adding a 7-year anchor point (Daloa only, group means) ---
# Data source: Donnees_semenciers_7ans.xlsx, sheet 'Pterocarpus' (n=12 individuals, 2 per group)
h7_individuals_m = [7.4, 9.5, 8, 7.4, 8, 10, 8, 5, 6, 7, 5, 8]  # meters, Daloa
h7_mean_cm = np.mean(h7_individuals_m) * 100
daloa_group_means = {6: 19.8, 12: 60.0, 24: 296.4}  # cm, from descriptive stats
t_ext = np.array([6, 12, 24, 84])
y_ext = np.array([daloa_group_means[6], daloa_group_means[12], daloa_group_means[24], h7_mean_cm])
for name, f, p0, k in [('linear', lin_f, [0, 10], 2), ('power', pow_f, [3, 1], 2),
                       ('exponential', exp_f, [15, 0.05], 2)]:
    popt, _ = curve_fit(f, t_ext, y_ext, p0=p0, maxfev=20000)
    rss = np.sum((y_ext - f(t_ext, *popt))**2)
    print(f"[7yr ext] {name}: AIC={aic_from_rss(rss, 4, k):.1f}")
try:
    popt, _ = curve_fit(gompertz, t_ext, y_ext, p0=[800, 5, 0.03], maxfev=50000,
                        bounds=([y_ext.max(), 0.5, 0.001], [5000, 50, 1]))
    rss = np.sum((y_ext - gompertz(t_ext, *popt))**2)
    print(f"[7yr ext] Gompertz: AIC={aic_from_rss(rss, 4, 3):.1f}, asymptote={popt[0]:.1f} cm")
except Exception as e:
    print("[7yr ext] Gompertz fit failed:", e)

# ---------------------------------------------------------------------
# 9. AXIS 5 - PHENOTYPIC PLASTICITY INDEX (RDPI, Figure 9)
# ---------------------------------------------------------------------

piv = df.pivot_table(index=['age_mois', 'groupe_semencier'], columns='localite',
                     values='long plant', aggfunc='mean').reset_index()
piv['plasticity'] = (piv['Daloa'] - piv['Korhogo']).abs() / (piv['Daloa'] + piv['Korhogo'])
mother_dbh = df[['groupe_semencier', 'DBH_semencier_cm']].drop_duplicates()
piv = piv.merge(mother_dbh, on='groupe_semencier', how='left')

sub_p = piv.dropna(subset=['plasticity', 'DBH_semencier_cm'])
r, p_val = stats.pearsonr(sub_p['DBH_semencier_cm'], sub_p['plasticity'])
print(f"Plasticity vs DBH: r={r:.3f} p={p_val:.3f} n={len(sub_p)}")

groups = [piv[piv.groupe_semencier == g]['plasticity'].dropna().values for g in range(1, 7)]
groups = [g for g in groups if len(g) > 0]
f, p_anova = stats.f_oneway(*groups)
print(f"Plasticity ANOVA by group: F={f:.3f} p={p_anova:.3f}")

# Plasticity vs age (increases with age)
sub = piv.dropna(subset=['plasticity'])
r_age, p_age = stats.pearsonr(sub['age_mois'], sub['plasticity'])
print(f"Plasticity vs age (linear): r={r_age:.3f} p={p_age:.4f}")
groups_age = [sub[sub.age_mois == a]['plasticity'].values for a in [6, 12, 24]]
f_age, p_age_anova = stats.f_oneway(*groups_age)
print(f"Plasticity ANOVA by age: F={f_age:.3f} p={p_age_anova:.4f}")

# ---------------------------------------------------------------------
# 10. AXIS 6 - ALLOMETRIC EQUATIONS (Tables 7-8, Figure 10)
# ---------------------------------------------------------------------

def fit_allom(y_col, x_col, label):
    s = df.dropna(subset=[y_col, x_col])
    s = s[(s[y_col] > 0) & (s[x_col] > 0)]
    X = sm.add_constant(np.log(s[x_col]))
    y = np.log(s[y_col])
    m = sm.OLS(y, X).fit()
    a = np.exp(m.params.iloc[0]); b = m.params.iloc[1]
```

```
    print(f"{label}: n={len(s)}, Y={a:.4f}*X^{b:.3f}, R2={m.rsquared:.3f}")

fit_allom('mas sc to plant', 'diam plant', 'Total biomass ~ diameter')
fit_allom('pd sc feuil', 'diam plant', 'Leaf biomass ~ diameter')
fit_allom('pd sc raci', 'diam plant', 'Root biomass ~ diameter')
fit_allom('surface feuil', 'long feuil', 'Leaf area ~ leaf length')

# Model comparison: power law vs quadratic log-log vs multiple (D+H)
sub = df.dropna(subset=['mas sc to plant', 'diam plant', 'long plant'])
sub = sub[(sub['mas sc to plant'] > 0) & (sub['diam plant'] > 0)]
logx = np.log(sub['diam plant']); logy = np.log(sub['mas sc to plant'])

X1 = sm.add_constant(logx)
m1 = sm.OLS(logy, X1).fit()
print(f"Power law: AIC={m1.aic:.1f}, R2={m1.rsquared:.3f}")

X2 = sm.add_constant(np.column_stack([logx, logx**2]))
m2 = sm.OLS(logy, X2).fit()
print(f"Quadratic log-log: AIC={m2.aic:.1f}, R2={m2.rsquared:.3f}")

X3 = sm.add_constant(np.column_stack([logx, np.log(sub['long plant'])]))
m3 = sm.OLS(logy, X3).fit()
print(f"Multiple (D+H): AIC={m3.aic:.1f}, R2={m3.rsquared:.3f}")
print(f"Equation: ln(B) = {m3.params.iloc[0]:.2f} + {m3.params.iloc[1]:.2f}*ln(D) + {m3.params.iloc[2]:.2f}*ln(H)")

# Root-to-shoot ratio (6 months only - see Data Quality Control)
sub_rs = df.dropna(subset=['pd sc raci', 'mas sc to plant'])
sub_rs = sub_rs[sub_rs['mas sc to plant'] > sub_rs['pd sc raci']]
sub_rs['ratio_RS'] = sub_rs['pd sc raci'] / (sub_rs['mas sc to plant'] - sub_rs['pd sc raci'])
print(sub_rs.groupby('age_mois')['ratio_RS'].agg(['count', 'mean', 'std']).round(3))

# ---------------------------------------------------------------------
# 11. CORRELATION MATRIX (Figure 11)
# ---------------------------------------------------------------------

corr_vars = ['long plant', 'diam plant', 'surface feuil', 'mas sc to plant', 'pd sc raci', 'long raci']
labels = ['Height', 'Collar diameter', 'Leaf area', 'Total biomass', 'Root biomass', 'Root length']
corr = df[corr_vars].corr()
corr.index = labels; corr.columns = labels
print(corr.round(2))

# ======================================================================
# End of script
# ======================================================================
```

S2. R script

The script below translates the Python pipeline into R, using lme4/lmerTest for the linear mixed models as stated in the manuscript's Methods section.

```
# ==================================================================
# Pterocarpus erinaceus - Complete analysis pipeline (R equivalent)
# UNVERIFIED BY THE AUTHORS - provided as a translation of the Python
# pipeline for readers preferring R / for cross-validation purposes.
# Please re-run and compare against the Python output before relying
# on these results for publication.
# ==================================================================
# Required packages: dplyr, tidyr, lme4, lmerTest, minpack.lm
# ==================================================================

library(dplyr)
library(tidyr)
library(lme4)
library(lmerTest)
library(minpack.lm)   # for nlsLM (robust nonlinear least squares)

# ------------------------------------------------------------------
# 1. DATA LOADING
# ------------------------------------------------------------------
# NOTE: the raw Excel files require substantial reshaping (one row per
# organ measurement -> one row per individual seedling). This step was
# performed in Python (pandas) for the published analysis; below we
# assume the cleaned master file 'pterocarpus_master_clean.csv' (produced
# by the Python pipeline) is used as the common starting point in R.

df <- read.csv("pterocarpus_master_clean.csv", stringsAsFactors = FALSE)
df$localite <- as.factor(df$localite)
df$groupe_semencier <- as.factor(df$groupe_semencier)
df$age_mois <- as.factor(df$age_mois)

mother_tree <- data.frame(
  groupe_semencier = 1:6,
  DBH_semencier_cm = c(12.4, 23.11, 28.41, 18.6, 25.94, 15.72),
  H_semencier_m    = c(8.31, 14.7, 17.65, 12.23, 13.59, 15.66),
  Masse_graine_moy_g = c(0.07, 0.08, 0.07, 0.07, 0.07, 0.07)
)

# ------------------------------------------------------------------
# 2. GERMINATION KINETICS AND CHI-SQUARE TEST (Figure 2)
# ------------------------------------------------------------------

n_total <- 138
germ_table <- matrix(c(97, 138 - 97,      # Daloa germinated / not germinated
                       85, 138 - 85),    # Korhogo germinated / not germinated
                     nrow = 2, byrow = TRUE,
                     dimnames = list(c("Daloa", "Korhogo"), c("Germinated", "Not")))
chisq.test(germ_table)

# Logistic model fit (nonlinear least squares)
days <- c(5, 10, 15, 20, 25, 30)
pct_daloa   <- c(18.1, 63.8, 82.6, 95.7, 100.0, 100.0)
pct_korhogo <- c(4.3, 42.0, 58.7, 66.7, 76.1, 76.1)

logistic_model <- function(t, Gmax, k, t50) Gmax / (1 + exp(-k * (t - t50)))

fit_daloa <- nlsLM(pct_daloa ~ logistic_model(days, Gmax, k, t50),
                   start = list(Gmax = 70, k = 0.3, t50 = 12))
fit_korhogo <- nlsLM(pct_korhogo ~ logistic_model(days, Gmax, k, t50),
```

```
                    start = list(Gmax = 70, k = 0.3, t50 = 12))
summary(fit_daloa)
summary(fit_korhogo)

# ---------------------------------------------------------------------
# 3. CLASSICAL ANOVA (site and age effects, Table 4)
# ---------------------------------------------------------------------

for (age in c(6, 12, 24)) {
  sub <- df[df$age_mois == age, ]
  for (col in c("long.plant", "diam.plant", "mas.sc.to.plant")) {
    if (col %in% names(sub)) {
      f <- aov(as.formula(paste(col, "~ localite")), data = sub)
      cat(age, col, "\n"); print(summary(f))
    }
  }
}

# ---------------------------------------------------------------------
# 4. AXIS 1 - CONTINUOUS REGRESSION (16 tests, Figure 5)
# ---------------------------------------------------------------------

for (age in c(6, 12, 24)) {
  sub_age <- df[df$age_mois == age, ]
  for (param in c("long.plant", "diam.plant", "mas.sc.to.plant")) {
    for (pred in c("DBH_semencier_cm", "H_semencier_m")) {
      sub2 <- sub_age[!is.na(sub_age[[param]]) & !is.na(sub_age[[pred]]), ]
      if (nrow(sub2) >= 5) {
        m <- lm(as.formula(paste(param, "~", pred)), data = sub2)
        cat(age, param, pred, "p=", summary(m)$coefficients[2, 4],
            "R2=", summary(m)$r.squared, "n=", nrow(sub2), "\n")
      }
    }
  }
}

# ---------------------------------------------------------------------
# 5. AXIS 2 - LINEAR MIXED MODELS WITH lme4/lmerTest (Table 5)
# ---------------------------------------------------------------------
# This is the analysis explicitly cited in the manuscript methodology
# ("Bates et al., 2015, lme4/lmerTest packages"). Please compare the
# p-values below against those reported in Table 5 and in the Python
# script's statsmodels output (section 6).

df_h <- df[!is.na(df$long.plant) & !is.na(df$DBH_semencier_cm) & !is.na(df$H_semencier_m) &
          !is.na(df$localite) & !is.na(df$age_mois) & !is.na(df$groupe_semencier), ]
model_height <- lmer(long.plant ~ DBH_semencier_cm * localite + H_semencier_m + age_mois +
                     (1 | groupe_semencier), data = df_h, REML = TRUE)
summary(model_height)
anova(model_height)

df_d <- df[!is.na(df$diam.plant) & !is.na(df$DBH_semencier_cm) & !is.na(df$H_semencier_m) &
          !is.na(df$localite) & !is.na(df$age_mois) & !is.na(df$groupe_semencier), ]
model_diam <- lmer(diam.plant ~ DBH_semencier_cm * localite + H_semencier_m + age_mois +
                   (1 | groupe_semencier), data = df_d, REML = TRUE)
summary(model_diam)
anova(model_diam)

df_b <- df[!is.na(df$mas.sc.to.plant) & df$mas.sc.to.plant > 0 &
          !is.na(df$DBH_semencier_cm) & !is.na(df$H_semencier_m) & !is.na(df$localite) &
          !is.na(df$age_mois) & !is.na(df$groupe_semencier), ]
df_b$log_biomass <- log(df_b$mas.sc.to.plant)
model_biomass <- lmer(log_biomass ~ DBH_semencier_cm * localite + H_semencier_m + age_mois +
```

```
                    (1 | groupe_semencier), data = df_b, REML = TRUE)
summary(model_biomass)
anova(model_biomass)

# ---------------------------------------------------------------------
# 6. AXIS 3 - MEDIATION ANALYSIS AND SOBEL TEST (Figure 6)
# ---------------------------------------------------------------------

germ_groups <- data.frame(
  groupe_semencier = 1:6,
  taux_germination = c(65.55, 66.66, 67.77, 71.99, 78.44, 77.50)
)
growth_by_group <- aggregate(long.plant ~ groupe_semencier, data = df, FUN = mean)
names(growth_by_group)[2] <- "mean_height"

path_df <- merge(mother_tree, germ_groups, by = "groupe_semencier")
path_df <- merge(path_df, growth_by_group, by = "groupe_semencier")

path_a <- lm(Masse_graine_moy_g ~ DBH_semencier_cm, data = path_df)
path_b <- lm(taux_germination ~ Masse_graine_moy_g, data = path_df)
path_c <- lm(mean_height ~ taux_germination, data = path_df)
path_total <- lm(mean_height ~ DBH_semencier_cm, data = path_df)

a <- coef(path_a)[2]; se_a <- summary(path_a)$coefficients[2, 2]
b <- coef(path_b)[2]; se_b <- summary(path_b)$coefficients[2, 2]

sobel_z <- (a * b) / sqrt(b^2 * se_a^2 + a^2 * se_b^2)
sobel_p <- 2 * (1 - pnorm(abs(sobel_z)))
cat("Sobel z =", sobel_z, " p =", sobel_p, "\n")

# ---------------------------------------------------------------------
# 7. AXIS 4 - NON-LINEAR GROWTH MODELS (Tables 6-7, Figures 7-8)
# ---------------------------------------------------------------------

aic_common_scale <- function(y, yhat, k) {
  n <- length(y)
  rss <- sum((y - yhat)^2)
  n * log(rss / n) + 2 * k
}

for (varcol in c("long.plant", "diam.plant")) {
  for (site in c("Daloa", "Korhogo")) {
    sub <- df[df$localite == site & !is.na(df[[varcol]]), ]
    t <- sub$age_mois %>% as.character() %>% as.numeric()
    y <- sub[[varcol]]

    m_lin <- lm(y ~ t)
    m_pow <- nlsLM(y ~ a * t^b, start = list(a = 3, b = 1.3))
    m_exp <- nlsLM(y ~ a * exp(b * t), start = list(a = 10, b = 0.1))

    cat(varcol, site, "- Linear AIC:", aic_common_scale(y, fitted(m_lin), 2), "\n")
    cat(varcol, site, "- Power AIC:", aic_common_scale(y, fitted(m_pow), 2), "\n")
    cat(varcol, site, "- Exponential AIC:", aic_common_scale(y, fitted(m_exp), 2), "\n")
  }
}

# Gompertz model (height), tested but not retained (poorly constrained asymptote)
gompertz <- function(t, A, B, k) A * exp(-B * exp(-k * t))
for (site in c("Daloa", "Korhogo")) {
  sub <- df[df$localite == site & !is.na(df$long.plant), ]
  t <- sub$age_mois %>% as.character() %>% as.numeric()
  y <- sub$long.plant
  tryCatch({
```

```
    m_gomp <- nlsLM(y ~ gompertz(t, A, B, k),
                    start = list(A = max(y) * 2, B = 5, k = 0.05),
                    lower = c(max(y), 0.1, 0.001),
                    upper = c(max(y) * 20, 100, 5))
    print(summary(m_gomp))
  }, error = function(e) cat(site, "Gompertz fit failed:", conditionMessage(e), "\n"))
}

# Exploratory 7-year extension (Daloa group means only, n=12 individuals)
t_ext <- c(6, 12, 24, 84)
y_ext <- c(19.8, 60.0, 296.4, 744.2)  # cm; 84-month value = mean of 12 individuals
m_gomp_ext <- nlsLM(y_ext ~ gompertz(t_ext, A, B, k),
                    start = list(A = 800, B = 5, k = 0.03),
                    lower = c(max(y_ext), 0.5, 0.001), upper = c(5000, 50, 1))
summary(m_gomp_ext)

# ---------------------------------------------------------------------
# 8. AXIS 5 - PHENOTYPIC PLASTICITY INDEX (RDPI, Figure 9)
# ---------------------------------------------------------------------

piv <- df %>%
  group_by(age_mois, groupe_semencier, localite) %>%
  summarise(mean_height = mean(long.plant, na.rm = TRUE), .groups = "drop") %>%
  pivot_wider(names_from = localite, values_from = mean_height)

piv$plasticity <- abs(piv$Daloa - piv$Korhogo) / (piv$Daloa + piv$Korhogo)
piv <- merge(piv, mother_tree[, c("groupe_semencier", "DBH_semencier_cm")], by = "groupe_semencier")

cor.test(piv$DBH_semencier_cm, piv$plasticity)
summary(aov(plasticity ~ as.factor(groupe_semencier), data = piv))
summary(aov(plasticity ~ age_mois, data = piv))
cor.test(as.numeric(as.character(piv$age_mois)), piv$plasticity)

# ---------------------------------------------------------------------
# 9. AXIS 6 - ALLOMETRIC EQUATIONS (Tables 7-8, Figure 10)
# ---------------------------------------------------------------------

sub <- df[!is.na(df$mas.sc.to.plant) & df$mas.sc.to.plant > 0 &
          !is.na(df$diam.plant) & df$diam.plant > 0, ]
logx <- log(sub$diam.plant)
logy <- log(sub$mas.sc.to.plant)

m_power <- lm(logy ~ logx)
m_quad  <- lm(logy ~ logx + I(logx^2))
m_multi <- lm(logy ~ logx + log(sub$long.plant))

AIC(m_power); AIC(m_quad); AIC(m_multi)
summary(m_multi)

# Root-to-shoot ratio (6 months only)
sub_rs <- df[!is.na(df$pd.sc.raci) & !is.na(df$mas.sc.to.plant) &
             df$mas.sc.to.plant > df$pd.sc.raci, ]
sub_rs$ratio_RS <- sub_rs$pd.sc.raci / (sub_rs$mas.sc.to.plant - sub_rs$pd.sc.raci)
aggregate(ratio_RS ~ age_mois, data = sub_rs, FUN = function(x) c(mean = mean(x), sd = sd(x)))

# ---------------------------------------------------------------------
# 10. CORRELATION MATRIX (Figure 11)
# ---------------------------------------------------------------------

corr_vars <- c("long.plant", "diam.plant", "surface.feuil", "mas.sc.to.plant",
               "pd.sc.raci", "long.raci")
cor(df[, corr_vars], use = "pairwise.complete.obs")
```

```
# =============================================================================
# End of script
# =============================================================================
```
